# Development and validation of versatile species-specific primer assays for eDNA monitoring and authentication of 10 commercially important Peruvian marine species

**DOI:** 10.1101/2024.10.22.619663

**Authors:** Alan Marín, Ruben Alfaro, Lorenzo E. Reyes-Flores, Claudia Ingar, Luis E. Santos-Rojas, Irina B. Alvarez-Jaque, Karen Rodríguez-Bernales, Cleila Carbajal, Angel Yon-Utrilla, Eliana Zelada-Mázmela

## Abstract

Molecular identification assays provide crucial support in the research and regulation of aquatic resources. Among them, species-specific primers provide strong discriminatory power for fast and simultaneous differentiation between closely related species. In this study, we used interspecific variations detected in two mitochondrial genes to develop species-specific primers for eDNA monitoring and identifying 10 fish and shellfish species commercially available within the Peruvian seafood sector. To ensure versatility and high specificity, our primers were subjected to PCR, qPCR, and sequencing methods, coupled with robust validation assays that included a) an in-silico stage using self-generated and public DNA sequences, b) an in-vitro stage using target species belonging to vouchered specimens, fresh and cooked commercial samples, early life stages, and broad taxa of non-target species, and c) an in-situ stage using eDNA samples from different Peruvian marine ecosystems. Our novel species-specific primers successfully passed the validation process with high efficiency and specificity in unequivocally identifying all target species with 100% accuracy and without cross-species amplifications, thereby making them valuable tools for eDNA monitoring, seafood authentication, and to identify and combat illegal, unreported and unregulated fishing. The identification assays presented herein can be used to support effective fishery management and conservation efforts not only in the Peruvian fishery sector but also in other countries where our target species also occur or are available as imported commodities.

## 1. Introduction

Accurate taxonomic identification is crucial in many domains including effective species management, biodiversity monitoring, food authentication, wildlife trafficking regulation, outbreak surveillance of pathogenic agents, and medical diagnosis [1–6], just to mention a few. DNA sequencing is undoubtedly one of the most reliable and accurate methods to achieve species-level identification. However, despite its high accuracy, DNA sequencing technologies are still time-consuming, labor-intensive, and relatively expensive [7, 8].

A cheaper and faster alternative molecular approach capable of accurately identifying species even from highly degraded DNA is based on species-specific primer (henceforth SSP) assays and Polymerase Chain Reaction (PCR). The application of SSPs for rapid species identification has raised extensive attention among researchers, certification bodies, and law enforcement agencies due to its accuracy, time and cost-effectiveness, practical ease, and simplicity of results interpretation [9, 10]. The specificity of SSPs relies on a perfect or near perfect complementarity between the primer and its target species’s template strand. Conversely, to reduce the potential for cross-species amplification, nucleotide mismatches must be present in the hybridization region of non-target species, especially within the primer’s 3’ end (the last 5 nucleotides of the 3’ end region). The most detrimental effects result from a mismatch present at the 3’-terminal (first nucleotide from the 3’ end), which disrupts polymerase activity, greatly diminishing PCR efficiency or leading to a complete abolishment of PCR amplification [11, 12].

SSPs are ideally designed to amplify specific short length (circa from 100 to 300 bp) mitochondrial alleles that are present in higher copy numbers per cell than nuclear genome copies [13, 14]. The characteristic small amplicon size yielded by SSPs is not only advantageous for the species identification of processed products by endpoint PCR [15–17], but it is also ideal to be used in qPCR platforms using different kinds of starting genetic material including genomic DNA from fresh or processed organic tissues [18–20], early planktonic life stages [21], homogenized tissues or non-invasive DNA obtained by mucus swabbing that can be amplified by direct qPCR without DNA isolation step [3, 5, 22], and different environmental DNA (eDNA) sources such as fresh and marine water environments [23–25], water from bags used for international fish transport [20], pond sediments [26], terrestrial soil [27], and airborne eDNA [28].

Species-specific eDNA detection is a fast, cost-effective, highly sensitive, and relatively new technique that relies upon the presence of free DNA molecules that have been shed by the organisms that inhabit a certain ecosystem. The genetic material is usually collected by filtration and PCR amplified using SSPs and tested for presence/absence of target species [29]. Currently, eDNA is globally and successfully used to determine the presence/absence of invasive, endangered, or commercially important aquatic species including fish, mollusks, crustaceans, amphibians, reptiles, and marine mammals [24, 30]. Furthermore, several species-specific eDNA studies reported strong correlations between target species abundance and eDNA concentrations of different aquatic organisms [31–35], thereby demonstrating its potential not only to accurately map the distribution of target species but also as a promising tool for assessing stock biomass fluctuations [36].

The Peruvian sea harbors a vast amount of biodiversity [37–39] of which at least 250 fish and 74 shellfish species interact with the artisanal fishery [40]. Fish and shellfish species from the families Centrolophidae (cojinovas), Haemulidae (grunts), Loliginidae (squids), Paralichthyidae (flounders), Pectinidae (scallops), and Serranidae (groupers, rock seabasses) are among the most demanded and highly exploited. However, despite its great seafood diversity, Peru still lacks an official list of standardized commercial names for seafood species and several studies have highlighted the high level of homonymy (one name used for two or more species: e.g. “*lenguado*” and “*mero*” are used to label many flatfish and grouper species respectively) and synonymy (two or more names used for a single species: e.g. “*mocosa*”, “*cojinova mocosa*”, and “*ojo de uva*” refer to the mocosa ruff *Schedophilus haedrichi*) in commercial names used within the fishery sector (38, 41–44). The abundant number of seafood species commercially available within the Peruvian fishery sector combined with the use of generic or ambiguous commercial names and weak monitoring mechanisms across the supply chain stages provide a fertile scenario for unintended mislabeling or intentional substitution of high-value species by cheaper ones [38, 45].

The first peer-reviewed article (and largest study to date) to reveal seafood mislabeling along different Peruvian regions (Tumbes, Lambayeque, La Libertad, Ancash, and Lima) throughout the seafood supply chain (from fish landing sites to wholesale markets, supermarkets, and restaurants) was published by our team in this journal [38]. In that study, Marín et al. [38] utilized a combination of full (COI, 16S rRNA, and D-Loop) and mini (COI and 12S rRNA) DNA barcode markers to authenticate a broad variety of national and imported seafood commodities including fish, crustacean, and mollusk species of different presentations including fresh (whole body, filets, fish roe), processed (cured, frozen filets, fish and shellfish burgers, instant noodles, canned), and commercial cooked, collected during a 21-month period (from July 2016 to March 2018) from 6 Peruvian regions, revealing that 26.72% of the analyzed samples collected in 5 regions were mislabeled. Subsequent peer-reviewed articles based on full DNA barcode (COI) analyses revealed higher mislabeling values within the Peruvian seafood sector. Thus, Biffi et al. [43] identified fish, squid, and a cetacean species (whole individuals, filets, cooked, and marinated) in samples collected between May and June 2017 from landing sites, wholesale and retail markets, supermarkets, and restaurants from Lima and Tumbes regions, revealing a 32.7% mislabeling rate. In another study based on COI barcodes by Velez-Zuazo et al. [46] it was reported a 43% mislabeling rate in fish samples (fresh, refrigerated, frozen, uncooked, and marinated) collected from September 2017 to February 2018 in wholesale fish markets, supermarkets, and restaurants from Lima region.

Based on the evidence reported in the aforementioned studies, it is by no means clear that seafood mislabeling has become a major concern within different levels of the Peruvian supply chain, affecting conservation efforts, effective fishery management, and consumer finances. Hence there is an urgent need for the development of molecular assays for rapid yet accurate identification of seafood species to combat mislabeling. Nevertheless, so far only 4 Peruvian marine species have been targeted for the development of SSPs including 3 fish [3, 18] and a bivalve species [15, 16]. Thus, given the widespread presence of mislabeling along the Peruvian supply chain and the scarcity of species-specific markers that can be used in different scenarios that require the detection of marine species, this study aimed to develop and evaluate the specificity, utility, and versatility of novel SSPs targeting 10 commercially important marine species. The novel SSP sets were validated across three different stages including 1) in-silico: using self-generated and public DNA sequences available from databases, 2) in-vitro: using hatchery-reared larvae, vouchered specimens, and forensic samples obtained from landing sites, fish markets, supermarkets, and restaurants, and 3) in-situ: using eDNA samples collected from northern and central Peruvian marine ecosystems.

## 2. Materials and methods

### 2.1 Case study

The present research is part of an extensive project that aimed to develop species-specific markers for fast identification of high-value Peruvian seafood species used for direct human consumption (DHC). A selected group of fish and shellfish species were selected as candidates for the development and evaluation of SSPs based on some of the following criteria: a) of high value or commercial interest, b) target of substitution by cheaper species or used to substitute other species, c) labeled with ambiguous commercial names: species whose commercial name is shared with two or more species, labeled with generic “umbrella” term, species commercialized with several names, and d) of similar morphological appearance such as congeneric or related species.

### 2.2 Biological samples collection and DNA extraction

#### 2.2.1 Fresh and cooked seafood samples collection and DNA isolation

Fresh samples of target and non-target species were collected from September 2017 to July 2024 from local commercial divers, fish landing sites, wholesale fish markets, local markets, supermarkets, and restaurants from different coastal Peruvian regions including Tumbes, Piura, Lambayeque, La Libertad, Ancash, and Tacna (Table 1 and S1 Table). For each target species, we collected a whole specimen that was used as a voucher for further morphological identification by a specialist taxonomist and molecular analysis by DNA barcoding. Voucher specimens were deposited in the DNA barcoding sample collection of the Laboratory of Genetics, Physiology, and Reproduction of the National University of Santa (Ancash, Peru). Small and medium-sized species were collected as whole individuals, while fin or muscle tissues were sampled from large-sized specimens. Photograph records of whole specimens were taken for all collected samples. In some instances, samples consisted of tissue samples or archived DNA from our previous research or kindly donated by colleagues. Additionally, commercially cooked samples of different presentations were collected from February 2018 to March 2024 in restaurants along the north-central Peruvian coast (Tumbes, Piura, Lambayeque, La Libertad, and Ancash regions). We targeted only seafood dishes whose labels covered the target species of this study: scallops, squids, flounders, rock seabasses, eye-grape seabass, mocosa ruff, groupers, and grunts. Tissues were rinsed with distilled water and preserved in 96% ethanol at −20 °C for further DNA analysis.

**Table 1.**
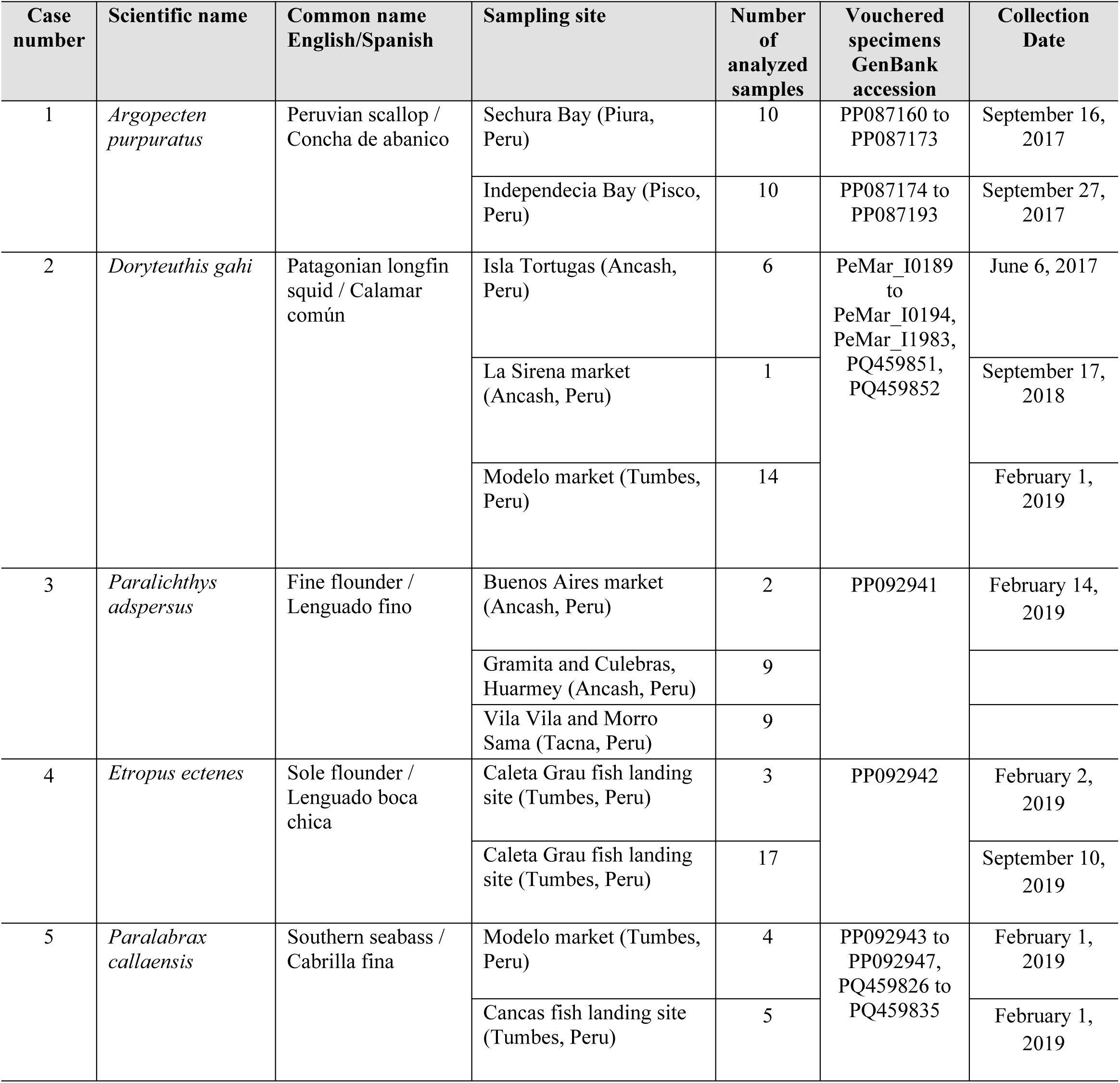
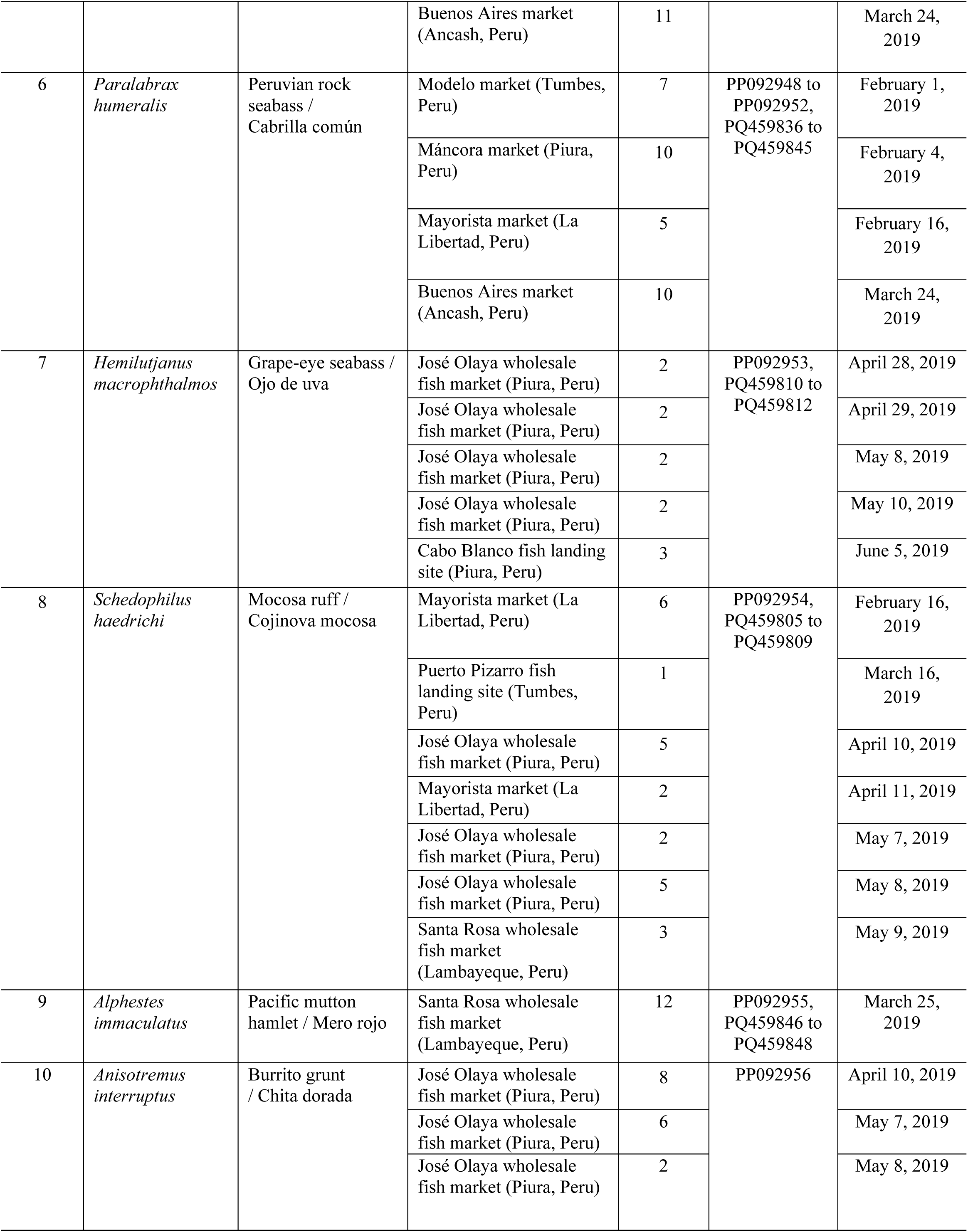
Target shellfish and fish species used for the validation assays of the species-specific primers developed in this study.

| Case number | Scientific name | Common name English/Spanish | Sampling site | Number of analyzed samples | Vouchered specimens GenBank accession | Collection Date |
| --- | --- | --- | --- | --- | --- | --- |
| 1 | *Argopecten purpuratus* | Peruvian scallop / Concha de abanico | Sechura Bay (Piura, Peru) | 10 | PP087160 to PP087173 | September 16, 2017 |
|  |  |  | Independencia Bay (Pisco, Peru) | 10 | PP087174 to PP087193 | September 27, 2017 |
| 2 | *Doryteuthis gahi* | Patagonian longfin squid / Calamar común | Isla Tortugas (Ancash, Peru) | 6 | PeMar_I0189 to PeMar_I0194, PeMar_I1983, PQ459851, PQ459852 | June 6, 2017 |
|  |  |  | La Sirena market (Ancash, Peru) | 1 |  | September 17, 2018 |
|  |  |  | Modelo market (Tumbes, Peru) | 14 |  | February 1, 2019 |
| 3 | *Paralichthys adspersus* | Fine flounder / Lenguado fino | Buenos Aires market (Ancash, Peru) | 2 | PP092941 | February 14, 2019 |
|  |  |  | Gramita and Culebras, Huarney (Ancash, Peru) | 9 |  |  |
|  |  |  | Vila Vila and Morro Sama (Tacna, Peru) | 9 |  |  |
| 4 | *Etropus ectenes* | Sole flounder / Lenguado boca chica | Caleta Grau fish landing site (Tumbes, Peru) | 3 | PP092942 | February 2, 2019 |
|  |  |  | Caleta Grau fish landing site (Tumbes, Peru) | 17 |  | September 10, 2019 |
| 5 | *Paralabrax callaensis* | Southern seabass / Cabrilla fina | Modelo market (Tumbes, Peru) | 4 | PP092943 to PP092947, PQ459826 to PQ459835 | February 1, 2019 |
|  |  |  | Cancas fish landing site (Tumbes, Peru) | 5 |  | February 1, 2019 |
|  |  |  | Buenos Aires market (Ancash, Peru) | 11 |  | March 24, 2019 |
| 6 | *Paralabrax humeralis* | Peruvian rock seabass / Cabrilla común | Modelo market (Tumbes, Peru) | 7 | PP092948 to PP092952, PQ459836 to PQ459845 | February 1, 2019 |
|  |  |  | Máncora market (Piura, Peru) | 10 |  | February 4, 2019 |
|  |  |  | Mayorista market (La Libertad, Peru) | 5 |  | February 16, 2019 |
|  |  |  | Buenos Aires market (Ancash, Peru) | 10 |  | March 24, 2019 |
| 7 | *Hemilutjanus macrophthalmos* | Grape-eye seabass / Ojo de uva | José Olaya wholesale fish market (Piura, Peru) | 2 | PP092953, PQ459810 to PQ459812 | April 28, 2019 |
|  |  |  | José Olaya wholesale fish market (Piura, Peru) | 2 |  | April 29, 2019 |
|  |  |  | José Olaya wholesale fish market (Piura, Peru) | 2 |  | May 8, 2019 |
|  |  |  | José Olaya wholesale fish market (Piura, Peru) | 2 |  | May 10, 2019 |
|  |  |  | Cabo Blanco fish landing site (Piura, Peru) | 3 |  | June 5, 2019 |
| 8 | *Schedophilus haedrichi* | Mocosa ruff / Cojinova mocosa | Mayorista market (La Libertad, Peru) | 6 | PP092954, PQ459805 to PQ459809 | February 16, 2019 |
|  |  |  | Puerto Pizarro fish landing site (Tumbes, Peru) | 1 |  | March 16, 2019 |
|  |  |  | José Olaya wholesale fish market (Piura, Peru) | 5 |  | April 10, 2019 |
|  |  |  | Mayorista market (La Libertad, Peru) | 2 |  | April 11, 2019 |
|  |  |  | José Olaya wholesale fish market (Piura, Peru) | 2 |  | May 7, 2019 |
|  |  |  | José Olaya wholesale fish market (Piura, Peru) | 5 |  | May 8, 2019 |
|  |  |  | Santa Rosa wholesale fish market (Lambayeque, Peru) | 3 |  | May 9, 2019 |
| 9 | *Alphestes immaculatus* | Pacific mutton hamlet / Mero rojo | Santa Rosa wholesale fish market (Lambayeque, Peru) | 12 | PP092955, PQ459846 to PQ459848 | March 25, 2019 |
| 10 | *Anisotremus interruptus* | Burrito grunt / Chita dorada | José Olaya wholesale fish market (Piura, Peru) | 8 | PP092956 | April 10, 2019 |
|  |  |  | José Olaya wholesale fish market (Piura, Peru) | 6 |  | May 7, 2019 |
|  |  |  | José Olaya wholesale fish market (Piura, Peru) | 2 |  | May 8, 2019 |

Genomic DNA from scallop samples was isolated from the adductor muscle using the automated DNA extractor iPrep TM (Invitrogen, Carlsbad, CA, USA). Genomic DNA from non-target bivalve, squid, and fish samples were isolated using the GeneJET Genomic DNA Purication Kit (Thermo Fisher Scientific, Carlsbad, CA, USA) or the standard phenol-chloroform protocol [48] from adductor muscle, tentacle tissues, or fin clips. DNA quantification was calculated using an Epoch spectrophotometer (BioTek Instruments, Winooski, VT, USA). Total gDNA was diluted to a final working concentration of 10-20 ng/μl and stored at −20 °C for further PCR analyses.

#### 2.2.2 Collection of early life stages of flounder and DNA extraction

To evaluate the performance of the SSPs in identifying fish species during early life stages, from July to September 2021 we collected premetamorphic (13 days after hatching, hereafter DAH), metamorphic larvae (31 DAH), and postmetamorphic juveniles (54 DAH) of *P. adspersus* from an aquaculture farm facility (Pacific Deep Frozen) located in Huarmey (Ancash, Peru). Larva and juvenile specimens were cold-anesthetized and preserved in RNA*later* solution (Invitrogen, Carlsbad, CA, USA). Genomic DNA was isolated from a small piece of fin clip using the standard phenol-chloroform protocol [47].

#### 2.2.3 eDNA collection, filtration of water samples, and eDNA extraction

In-situ testing should include eDNA samples from environments where the target species is present and also from environments where the target species is absent [13]. Therefore, we collected water samples in 5 field surveys from 7 different inshore stations belonging to two coastal bays with cultured and wild *A. purpuratus* stocks (Sechura Bay in Piura region and Samanco Bay in Ancash region), a bay with only wild *A. purpuratus* (Tortugas Bay in Ancash region), and an offshore station out of the distribution range of *A. purpuratus* populations (La Cruz, Tumbes region) (Fig 1). The first field survey was performed in October 2017 at Parachique station (Sechura Bay) using 0.5 L new plastic containers and nitrocellulose filter membranes (0.45 μm pore size and 47 mm diameter), eDNA samples were collected from the bottom (5 to 6 m depth), midwater (2-3 m depth), and surface (0.5 m depth). The second field survey was carried out in February 2019 at Samanco Bay (Ancash) using 1 L sterilized laboratory glass bottles and nitrocellulose filter membranes (0.45 μm pore size and 47 mm diameter), eDNA samples were collected from the surface (0.5 m depth) water column. Samples from the third and fourth field surveys consisted of repurposed marine eDNA filters originally collected for a metabarcoding study targeting sponge-associated bacterial communities, samplings were carried out from April to May 2022 at Bayóvar and Matacaballo stations from Sechura Bay (Piura region) and three stations from Tortugas Bay (Ancash region) using 1 L sterilized laboratory glass bottles and mixed cellulose ester (MCE) filter membranes (0.45 μm pore size and 47 mm diameter), eDNA samples were collected from the bottom water column (3 to 6 m depth, depending on each station) in close proximity to sponge communities. The field survey in the offshore location namely La Cruz (Tumbes region) was performed in December 2017. eDNA samples were collected from the surface (0.5 m depth) water column using 1 L sterilized laboratory glass bottles and nitrocellulose filter membranes (0.45 μm pore size and 47 mm diameter). Seawater samples were filtered within 45 min of collection, except for the sample collected at La Cruz offshore station which was kept in a cooler box containing frozen gel packs and filtered after four hours of collection. All eDNA samples were filtered using a manual vacuum pump connected to a 500 ml magnetic filter funnel (Rocker, Rocker Scientific Company Limited, Taiwan). To avoid contamination during field samplings, all materials (cooler box, gel packs, water containers, tweezers) were soaked in 25% bleach for 30 min, rinsed with 70% ethanol and distilled water, and sterilized under UV light. Disposable nitrile gloves were used during all filtering steps and replaced by new ones before filtering a new eDNA sample. After each round of filtration and before filtering the next sampling station, all equipment including the filtering funnel were soaked in 25% bleach for at least 10 min and rinsed with 70% ethanol and distilled water. For each sampling site, blank control samples were obtained by filtering 1 L of distilled water. After the filtration was completed, filters were carefully folded using sterilized tweezers and stored in 2 ml microtubes containing 96% ethanol. All filters were transported to the laboratory in a cooler box containing frozen gel packs. Once at lab, filters were immediately kept at −20 °C until DNA extraction.

**Fig 1.**
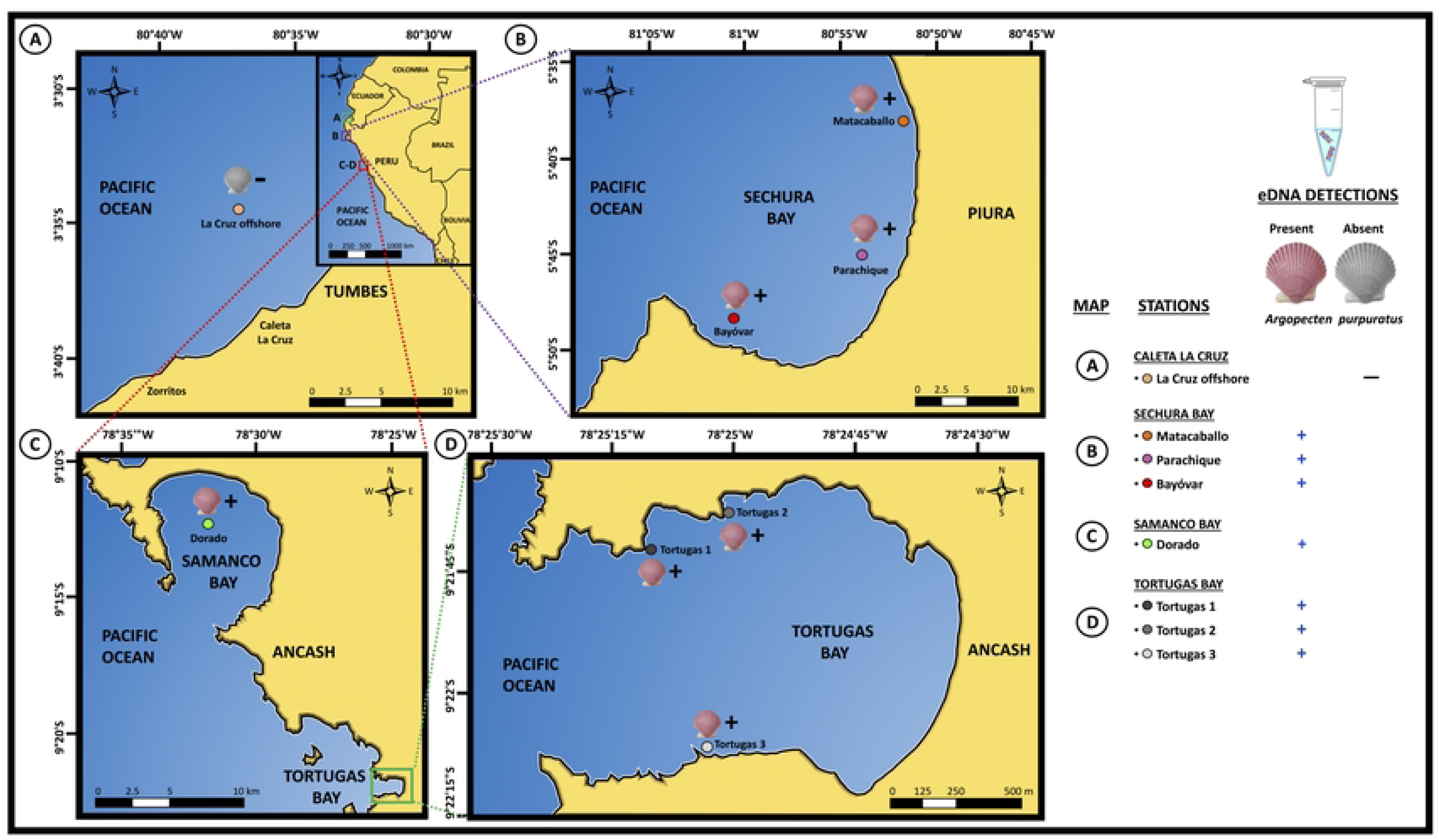
Map of the Peruvian eDNA sampling stations and eDNA detection results of *Argopecten puprpuatus*. Map A: Caleta La Cruz sampling station (Tumbes region). Map B: Sechura Bay (Piura region) showing Matacaballo, Parachique, and Bayóvar sampling stations. C: Samanco Bay (Ancash region) showing El Dorado sampling station. Map D: Tortugas Bay (Ancash region) showing three sampling stations namely Tortugas 1, Tortugas 2, and Tortugas 3. Positive detections of *A. purpuratus* eDNA are depicted by a “+” sign and a colored scallop valve, negative detections of *A. purpuratus* eDNA are depicted by a “-” sign and a scallop valve in black and white.

All eDNA extractions (eDNA-containing filters and blank control samples) were performed in a clean laboratory completely separated from pre-PCR and PCR rooms, following strict decontamination procedures (disinfection with 10% bleach, 70% ethanol, and UV-light sterilization) and use of disposable materials. In order to prevent contamination, only aerosol barrier tips were used during eDNA extractions. Filter membranes were removed from the ethanol, cut in half, air-dried, and shredded into small pieces using a sterile scalpel blade. Only eDNA extractions from Tortugas 1 and Tortugas 3 stations were performed using half and also quarter filters. Briefly, each shredded half (or quarter) filter was placed in a 2 ml Eppendorf tube and 600 μl lysis buffer (10 mM Tris-HCl, 5 mM EDTA, 20 mM NaCl, 1% SDS) and 50 μl proteinase K (10 mg/ml) were added to each filter tube and incubated at 55°C for 3 hours in a shaking incubator, tubes were vortexed every 15 min. Then, 600 μl phenol was added and mixed by inversion and incubated for 15 min at room temperature in a shaker (600 rpm). After that, 600 μl chloroform was added and mixed by inversion and incubated for 20 min at room temperature in a shaker (600 rpm). Samples were centrifuged at 13500 rpm at room temperature for 15 min. To precipitate eDNA, 500 μl of the aqueous phase was transferred to a clean 2 ml tube, mixed with 50 μl of 3M sodium acetate (NaOAc) and 1250 μl of 99% cold ethanol and incubated at −20 °C overnight. Next, the microtube was centrifuged at 13500 rpm for 30 min at 4 °C, and the ethanol was poured out. The pellet was washed with 500 μl of 70% ethanol, centrifuging at 13500 × rpm for 5 min at 4 °C, the ethanol was removed and the pellet was dried at 37 °C for 7 min. Once dried, the pellet was dissolved in 50 μl 1× TE Buffer pH 8.0 (Thermo Fisher Scientific Baltics, Vilnius, Lithuania).

### 2.3 Species authentication of voucher specimens and non-target species by DNA sequence analysis

The species identity of 34 *A. purpuratus* specimens was verified based on the DNA sequence of a partial fragment of the mitochondrial 16S rRNA (670 bp) amplified using a common *Argopecten* primer PurvenF developed herein and the universal primer 16Sbr [48] (S2 Table) under the following thermal cycling conditions: initial denaturation at 94 °C for 5 min, followed by 30 cycles of 94 °C for 20 s, 60 °C for 20 s, and 72 °C for 25 s, and a final extension step at 72 °C for 7 min. PCR reaction mixtures consisted of a 20 μl final volume containing 0.1 μl of Maximo *Taq* DNA Polymerase (GeneOn, GmbH, Nurnberg, Germany), 2.5 μl buffer 10×, 1 μl dNTPS (2.5 mM), 0.08 μl each primer (50 μM), 0.38 μl MgCl_2_ (100 μM), 1 μl of template DNA (10 ng/μl), and 14.86 μl ultrapure H_2_O. Successful PCR amplification products were visualized in a 1.5% agarose gel (EMD Millipore, Billerica, MA, USA) electrophoresis, amplicons were stained with GelRed Nucleic Acid Gel Stain (BIOTREND Chemikalien GmbH, Köln, Germany). PCR products were Sanger sequenced in both directions at Macrogen (Seoul, South Korea). All obtained scallop sequences were deposited in GenBank/DDBJ/EMBL DNA databases with accession numbers from PP087160 to PP087193.

Fish and squid voucher specimens of target and non-target species were morphologically identified by an expert taxonomist and by DNA barcoding assay using the universal primer sets 16Sar-L/16Sbr-H [48], FishF1/FishR1 [49], or LCO1490/HCO2198 [50] (S2 Table) with amplification conditions described by Marín et al. [38]. PCR products were purified using FastAP Thermosensitive Alkaline Phosphatase and Exonuclease I (Thermo Fisher Scientific, USA) following the manufacturer’s instructions and sequenced using the same primer set on an ABI PRISM 3500 Genetic Analyzer (Applied Biosystems, Hitachi, Foster City, CA, USA) using the BigDye terminator v3.1 Cycle Sequencing Kit (Applied Biosystems, Waltham, Massachusetts, USA) at the Laboratory of Genetics, Physiology, and Reproduction (National University of Santa, Peru). All obtained sequences were deposited in GenBank/DDBJ/EMBL DNA databases with accession numbers from PP092941 to PP092966 and PQ459826 to PQ459855.

Identification of all DNA sequences at the species level was accomplished by using both the Barcode of Life Data System (BOLD, http://www.boldsystems.org) selecting “species level barcode records” database and Basic Local Alignment Search Tool (BLAST) on the National Center for Biotechnology Information (NCBI, http://www.blast.ncbi.nlm.nih.gov/Blast.cgi). Current accepted scientific names were checked in The World Register of Marine Species (WORMS, available at http://www.marinespecies.org).

### 2.4 In-silico analysis for SSP design

Target mitochondrial genes for the designing of SSPs were selected based on high interspecific variability reported in fish and shellfish identification studies [15, 38, 51, 52]. Thus, three mitochondrial genes were selected: *cytochrome oxidase subunit I* (COI), *16S ribosomal RNA* (16S rRNA), and *12S ribosomal RNA* (12S rRNA). Aiming to include as many DNA sequences as possible from target and non-target species, we used self-generated and reference sequences retrieved from BOLD and GenBank databases. All sequences were multialigned in MEGA 7 software [53] generating different multi-sequence matrices that were used for intra and interspecific variation analyses of the putative SSP. At least three SSP sets for each target species were designed using AlleleID 7.0 software (PREMIER Biosoft, USA) or manually by searching non-conserved homolog regions at the 3’ end (defined as the last 5 nucleotides from the primer’s 3’ end [11]) of the primer hybridization site of closely related (e.g. congeneric) and non-related non-target species to ensure the specific amplification only in target species.

All SSP sets were designed to yield amplicon lengths of between 100 and 300 bp and to work at high annealing temperatures (≥ 60 °C) to ensure reaction astringency. Candidate oligos were tested in-silico for secondary structure formation (hairpins, homo and hetero-dimers) using the IDT OligoAnalyzer tool (available at <http://www.idtdna.com>) and The Sequence Manipulation Suite [54] (available at <http://www.bioinformatics.org/sms2/index.html>). Lastly, the specificity of the candidate oligos was analyzed in the NCBI GenBank database using Primer-BLAST [55] selecting “nr” non-redundant nucleotide database and organisms limited to Class Bivalvia (taxid:6544) in scallop primer analyses, the orders Myopsida (taxid:551347) and Oegopsida (taxid:34542) in squid primer analyses, and the Parvphylums Osteichthyes (taxid:7898) and Chondrichthyes (taxid:7777) in fish primer analyses. Any match to freshwater species and species not found in the Eastern Pacific were disregarded. All designed oligonucleotides were synthesized by Integrated DNA Technologies (IDT, Coralville, Iowa, USA) and Invitrogen (Carlsbad, CA, USA).

### 2.5 In-vitro SSP validation assays by endpoint PCR and qPCR

#### 2.5.1 Optimization and validation of endpoint PCR assay

To evaluate the species specificity of each SSP set, we first determined the maximum optimal annealing temperature by performing a gradient annealing temperature analysis ranging from 60 to 64 °C. To determine the minimum primer concentration that resulted in a reliable and specific PCR product, concentrations between 100 and 500 nM were evaluated at 50 nM increments. The optimal annealing temperature and primer concentration were initially tested in 5 individuals of the target species. All PCR reactions were performed in a Veriti 96 Well thermal cycler (Applied Biosystems, Foster City, CA, USA) using Maximo *Taq* DNA Polymerase 2x-preMix (GeneOn GmbH, Nurnberg, Germany), which include ammonium sulfate ((NH_4_)_2_SO_4_) on its buffer system. The NH_4_ ions destabilize weak hydrogen bonds and mismatched bases present in non-target loci and primer dimers enhancing the yield of specific PCR products [56]. The specificity of the newly designed primers was tested in-vitro using DNA extracted from all collected individuals of each target species (ranging from 11 to 32 specimens), where we expected to obtain a single PCR product of the expected size.

SSP validation assays for target shellfish species were performed against different non-target bivalve (7 species from 6 families) and squid (4 species from 2 families) species (S1 Table). The yield of artifacts resulting from the interaction between the SSP and endogenous control primer sets during the duplex PCR trials prevented us from including an internal control reaction during the specificity validation assays targeting the Peruvian scallop and the common squid against non-target species. We discarded the possibility of the presence of inhibitors or poor quality of non-target species’ DNAs because during species authentication analyses using the DNA barcoding approach we obtained abundant PCR products. Therefore, the specificity validation assays of *A. purpuratus* and *D. gahi* were based on the total absence of PCR products in non-target species.

For SSPs validation against non-target fish species, a duplex PCR was standardized using the SSP set (COI or 16S rRNA gene, depending on species) in a single reaction with an internal endogenous control using the fish universal primers MiFish-U [51] (S2 Table) designed to amplify a small fragment (circa 220 bp) of the mitochondrial 12S rRNA gene. Thus, in the absence of target DNA, non-target fish species (27 fish species from 19 families, S1 Table) were expected to yield only the universal 12S rRNA gene amplicon, resulting in the visualization of a single band in the agarose gel, while target species yielded two PCR products corresponding to the SSPs and the endogenous control bands. When possible, in order to validate the high specificity of our SSPs and to evaluate if they can be used to distinguish between closely related species, up to 22 individuals of non-target congeneric species were included in the PCR validation trials. PCR reactions were performed in a Veriti 96 Well thermal cycler (Applied Biosystems, Foster City, CA, USA) using Maximo *Taq* DNA Polymerase 2×-preMix (GeneOn GmbH, Nurnberg, Germany). All PCR reactions were visualized using a 1.5% agarose gel (EMD Millipore, Billerica, MA, USA) electrophoresis, amplicons were stained with GelRed Nucleic Acid Gel Stain (BIOTREND Chemikalien GmbH, Köln, Germany).

#### 2.5.2 Optimization and validation of qPCR assays

The specificity and efficiency of the SSP set for the identification of the target scallop species (*A. purpuratus*) was additionally evaluated in qPCR assays performed on a LightCycler 480 II (Roche Diagnostics GmbH, Penzberg, Germany) and run in triplicate on 96-well reaction plates (Roche Diagnostics GmbH, Mannheim, Germany) following MIQE guidelines [57]. qPCR mix reactions and cycling protocols for the identification of *A. purpuratus* are shown in Fig 3. The 2x SYBR Green I Master (Roche Diagnostics, Mannheim, Germany) kit was used in all experiments. A melting curve analysis was conducted from 60 °C for 1 min with a rate of 2.2 °C per second up to 95 °C with a continuous acquisition. All reactions were performed using a designated set of micropipettes for qPCR use only. A positive control (20 ng of target species gDNA) and a no-template control (NTC: 2 μl ultrapure H_2_O instead of template DNA) reaction were included in each qPCR run. Primer specificity confirmation was checked by melting curve analysis showing single sharp peaks only in target species with no visualization of secondary peaks with lower melting temperature (primer dimers).

The standard curve method was used to determine the qPCR reaction efficiency and the limit of detection (LOD), which was defined as the lowest amount of target DNA in a sample that can be reliably detected with > 95% amplification success [57]. Standard curves were constructed using 17 ng of gDNA from *A. purpuratus* that was obtained using a commercial DNA extraction kit based on magnetic beads technology (iPrep TM purification instrument, Invitrogen, Carlsbad, CA, USA). The gDNA was 10-fold serially diluted from 17 ng to 1.7 × 10^-8^ ng, and subjected to qPCR to construct the standard curves, using 2 μl of each standard and three technical replicates for each dilution. qPCR was performed using the qPCR mix reactions and cycling protocols shown in Fig 3, with the only modification being that total amplification cycles were 45 instead of 40. qPCR efficiency was calculated by plotting the Ct values of the dilution series against the logarithm of the DNA concentration from each standard dilution. The quantification cycles (Ct values) and amplification efficiency were calculated using the LightCycler 480 II Software (version 1.5.1.62) under the “Abs Quant/2nd Derivative Max” analyses, from the slope of the linear regression using the equation E = 10^(− 1/slope)^. Mean values, standard deviations, standard curve figures, and *R^2^* values were obtained with Excel 2016 for macOS.

### 2.6 Direct endpoint PCR and qPCR assays in commercial scallop samples

To further evaluate the performance of the *A. purpuratus* SSP set ARGOF/ARPU129R as a potential field deployable tool, we performed a direct qPCR assay (without DNA isolation step) using genetic material collected by non-invasive sampling of 5 fresh and 5 frozen individuals of *A. purpuratus*, bought in a local market and a supermarket, respectively. The non-invasive sampling was performed by gently rubbing a sterile cotton swab against the scallops’ adductor muscles (panel A in S2 Fig). Swabs used in fresh and frozen scallops were immediately placed in 1.5 ml Eppendorf tubes containing 1400 μl and 400 μl of 1× phosphate buffer saline (PBS), respectively. Before endpoint PCR and qPCR amplifications, the Eppendorf tubes were vortexed, spun down, and incubated for 10 min at 55 °C. Direct endpoint PCR and qPCR reactions were performed following the same PCR and qPCR amplification protocols described in Fig 3.

### 2.7 In-situ SSP validation assays

#### 2.7.1 In-situ validation assay of eDNA samples by qPCR

The reliability and specificity of the SSP sets targeting *A. purpuratus* were further tested to ascertain its presence or absence in field environmental samples. qPCR amplifications were conducted in triplicate in a LightCycler 480 II (Roche Diagnostics GmbH, Penzberg, Germany), the 2x SYBR Green I Master (Roche Diagnostics, Mannheim, Germany) kit was used in all qPCR eDNA runs on 96-well reaction plates (Roche Diagnostics GmbH, Mannheim, Germany) with the parameters shown in Fig 3, with two main modifications: 1 μl of each primer (10 μM) was used making a final SSP concentration of 0.5 μM in each reaction and 45 amplification cycles were used instead of 40. Environmental DNA amplifications were considered as positive detections only if at least two out of three qPCR technical replicates displayed fluorescent signal above the threshold in one field sample [58] and showed 100% sequence identity with target species after Sanger sequencing.

To quantify the DNA concentration present in each environmental sample, a standard curve was constructed using gDNA from *A. purpuratus* extracted with magnetic beads technology (iPrep TM purification instrument, Invitrogen, Carlsbad, CA, USA) in 10-fold serial dilutions ranging from 10 ng to 1.0 × 10^-11^ ng. qPCR conditions were the same as described in the previous paragraph. The standard curve method was also used to determine the limit of detection (LOD), which was defined as the lowest concentration of the standard dilutions that gave at least one positive amplification out of the 3 replicates. This LOD definition is in accord with previous eDNA published articles based on species-specific assays of different aquatic organisms [59–61].

#### 2.7.2 Inhibition testing

To determine the presence of inhibitors in the environmental samples that can lead to potential false negative results, all eDNA samples and negative field controls were spiked with an exogenous internal positive control (IPC) obtained from a freshwater fish species ensuring the complete absence of the exogenous DNA control in the marine environmental samples. The IPC consisted of 1 μl of 5 ng/μl of gDNA of a male specimen of the Amazonian giant fish *Arapaima gigas* (*paiche*) that was qPCR-amplified using the primer set MSR_129 previously developed for the specific genotypic sexing of the *paiche* [62]. eDNA samples were deemed inhibited if qPCR resulted in a Ct difference > 2 between the eDNA samples and the positive control of pure *paiche* DNA [25]. The IPC qPCR assay was performed in duplicate using 10 μl of 2x SYBR Green I Master (Roche Diagnostics, Mannheim, Germany), 0.2 μl each MSR_129 primer, 1 μl of *paiche* gDNA (5 ng/μl), and either 2 μl of eDNA template or 2 μl ultrapure H_2_O for the no-template control, in a 20 μl total reaction volume. The qPCR amplification parameters were identical to those described by López-Landavery et al. [62].

### 2.8 Authentication of positive eDNA amplicons by Sanger sequencing

In order to verify the species identity of the positive eDNA amplifications, positive qPCR amplicons (one per each positive replicate) were Sanger sequenced bidirectionally using the same SSP set ARGO/ARPU129R. Forward and reverse electropherograms were trimmed and aligned to obtain consensus sequences that were contrasted against the complete mitochondrial genome of *A. purpuratus* (GenBank accession KF601246).

### 2.9 Applicability of SPPs in commercial cooked fish samples from restaurants

Aiming to evaluate the applicability of our SSP sets to authenticate commercial cooked samples collected from restaurants, different cooked seafood presentations (cebiche, steamed, fried, seafood rice, and mixed fried seafood) were tested with our SSP assays (Table 4). Collected cooked samples included scallops, squids, rock seabasses, grape-eye seabasses, and grunts. For the identification of rock and grape-eye seabass labeled samples, two different duplex PCRs were performed. A first duplex PCR for “*cabrillas*” (rock seabasses) consisted of SSP sets PACA163F/R (targeting *P. callaensis*) and PAHU288F/R (targeting *P. humeralis*) in a single reaction tube using the PCR cocktail and amplification conditions shown in Panel A of S3 Fig. Results interpretation was based on the visualization of a single PCR product of 163 bp or 288 bp in length in each reaction, indicating the presence of DNA from *P. callaensis* or *P. humeralis*, respectively. The species identity of the PCR products from cooked “*cabrilla*” samples was confirmed by bidirectional Sanger sequencing using the same SSPs. For the identification of samples labeled as “*ojo de uva*”, which may refer either to the grape-eye seabass *H. macrophthalmos* or the mocosa ruff *S. haedrichi*, we performed a duplex PCR including the SSP sets HEMA122F/R and SCHA244F/R, under the amplification conditions shown in Panel B of S3 Fig. The correct identification of both species in a single PCR reaction was possible given the different amplicon sizes of 122 bp for *H. macrophthalmos* and 244 bp for *S. haedrichi*. All PCR results were verified by 1.5% agarose gel (EMD Millipore, Billerica, MA, USA) electrophoresis.

## 3. Results

Based on the criteria mentioned in the Materials and Methods section, different species belonging to the following shellfish and fish groups were selected as target species for SSP design: 1) scallops, 2) squids, 3) flounders, 4) rock seabasses, 5) seabasses, 6) cojinovas, 7) groupers, and 8) grunts. Among the different putative SSPs evaluated in several species belonging to the abovementioned seafood groups, successful SSP assays (those that did not show unspecific or cross-species amplifications) were optimized and validated using 201 specimens belonging to 10 target species (Fig 2, Table 2) whose natural distribution range from Central to South East Pacific and South West Atlantic.

**Fig 2.**
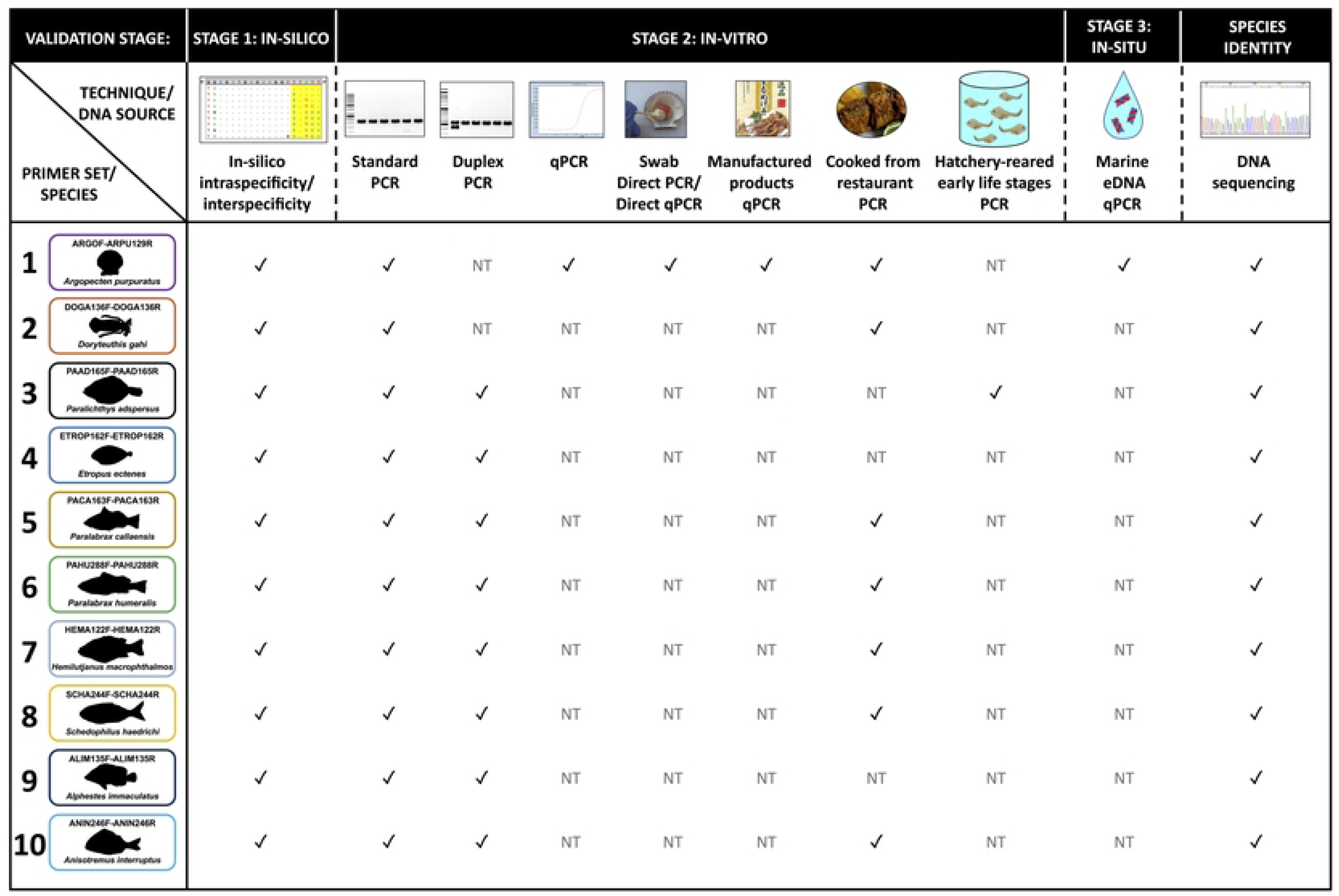
PCR analysis types and validation stages used for each species-specific primer sets. NT: not tested.

**Table 2.**
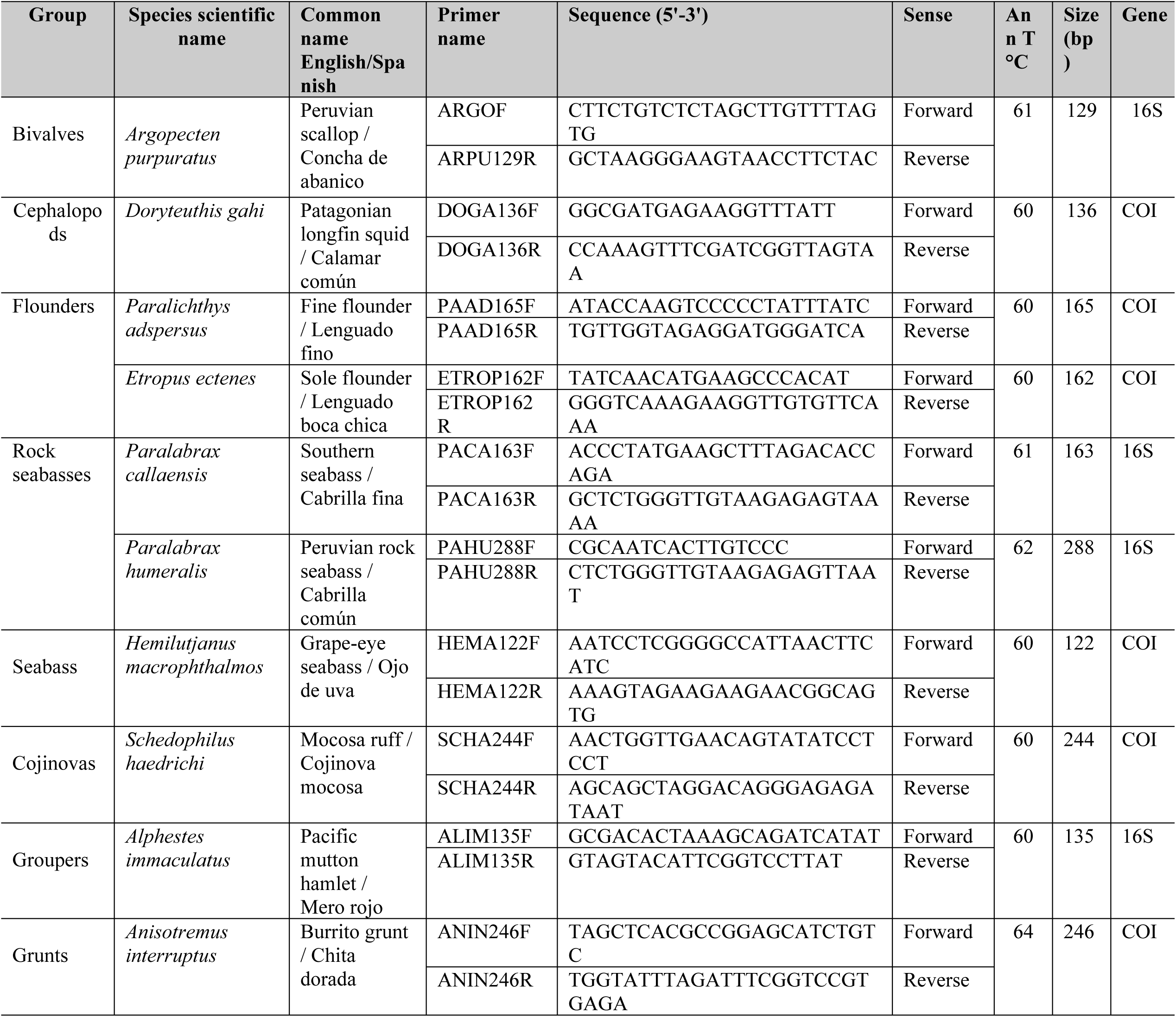
Species-specific primer sequences designed for the identification of 10 Peruvian marine species.

The shellfish species group with validated molecular identification assays was represented by the Peruvian scallop *Argopecten purpuratus* (Fig 3) and the Patagonian longfin squid *Doryteuthis gahi* (Fig 4). The fish species group included the fine flounder *Paralichthys adspersus* (Fig 5) and the sole flounder *Etropus ectenes* (Fig 6), the southern rock bass *Paralabrax callaensis* (Fig 7) and the Peruvian rock seabass *P. humeralis* (Fig 8), the grape-eye seabass *Hemilutjanus macrophthalmos* (Fig 9), the mocosa ruff *Schedophilus haedrichi* (Fig 10), the Pacific mutton hamlet *Alphestes immaculatus* (Fig 11), and the burrito grunt *Anisotremus interruptus* (Fig 12). PCR mix protocols and amplification conditions used in the validation of all SSPs are shown in Fig 3 to Fig 12.

**Fig 3.**
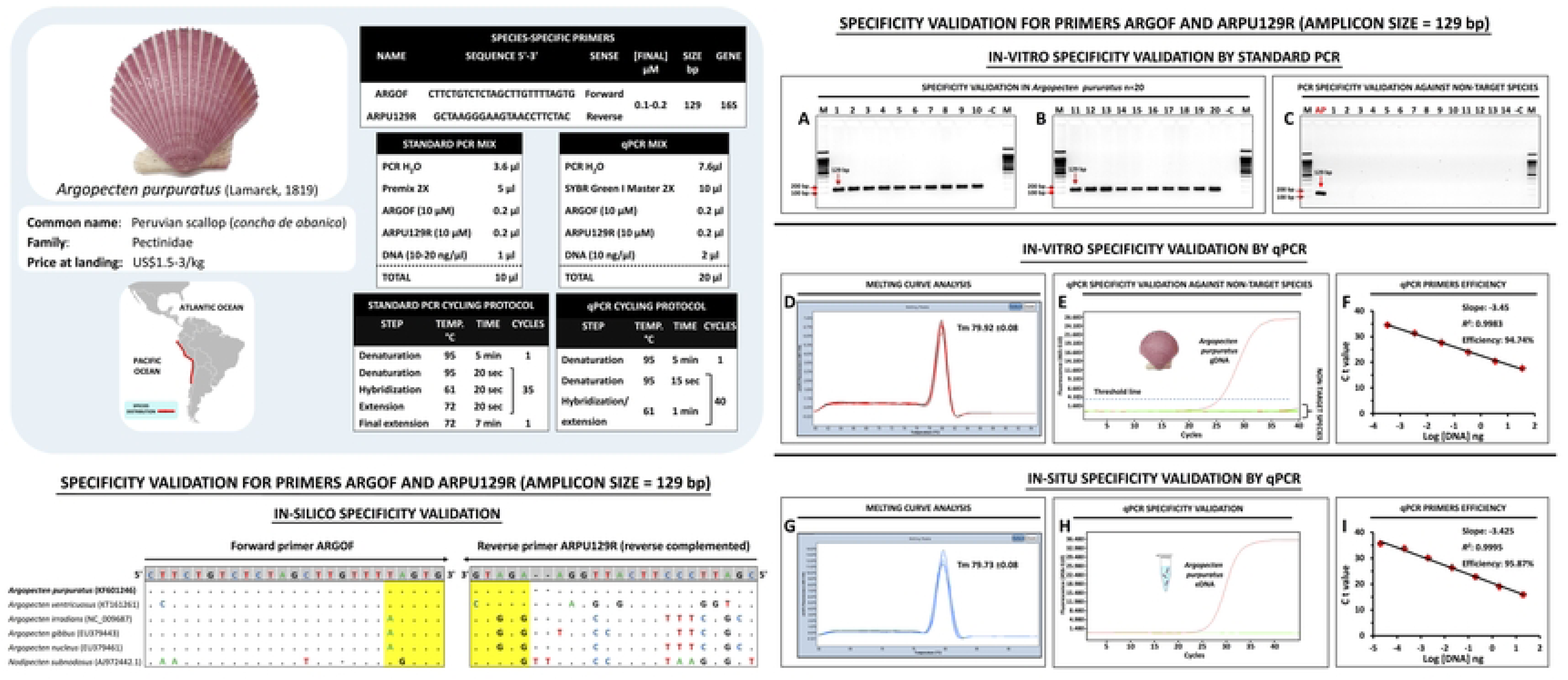
Standardized species-specific primer protocols and validation assay results (in-silico, in-vitro, in-situ) for the identification of *Argopecten purpuratus*. In the in-silico validation panel, the nucleotides highlighted in yellow correspond to the 3’ ends of each primer hybridization region. Panels A and B: PCR electrophoresis results of the primer set ARGOF/ARPU129R showing the positive detection (amplicon size 129 bp) in 20 individuals of *A. purpuratus*. Panel C: PCR electrophoresis results of the primer specificity validation against non-target bivalve species, well “AP” is the positive control (*A. purpuratus*), wells 1 to 8: *A. ventricosus*, well 9: *Pteria sterna*, well 10: *Striostrea prismatica*, well 11: *Atrina maura*, well 12: *Perumytilus purpuratus*, well 13: *Aulacomya atra*, well 14: *Gari solida*. In-vitro specificity validation by qPCR analysis of fresh individuals of *A. purpuratus* displayed in panel D: melting curve analysis result, panel E: specificity validation against non-target bivalve species (*A. maura*, *A. atra*, *A. ventricosus*, *G. solida*, *P. purpuratus*, *P. sterna*, and *S. prismatica*), panel F: qPCR efficiency results obtained by the standard curve method. In-situ specificity validation by qPCR analysis of *A. purpuratus* eDNA displayed in panel G: melting curve analysis result, panel H: specific detection of *A. purpuratus* in eDNA samples, panel F: eDNA-qPCR efficiency results obtained by the standard curve method.

**Fig 4.**
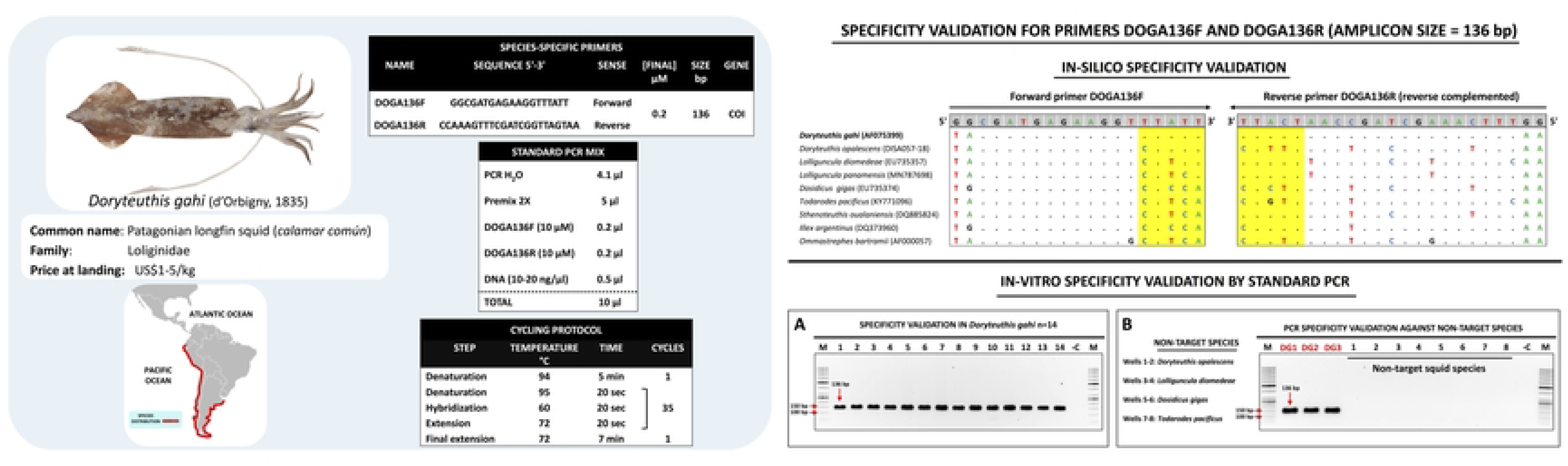
Standardized species-specific primer protocols and validation assay results (in-silico and in-vitro) for the identification of *Doryteuthis gahi*. In the in-silico validation panel, the nucleotides highlighted in yellow correspond to the 3’ ends of each primer hybridization region. Panel A: PCR electrophoresis results of the primer set DOGA136F/R showing the positive detection (amplicon size 136 bp) in 14 individuals of *D. gahi*. Panel B: PCR electrophoresis results of the primer specificity validation against non-target squid species, wells “DG1” to “DG3” are positive controls for *D. gahi*.

**Fig 5.**
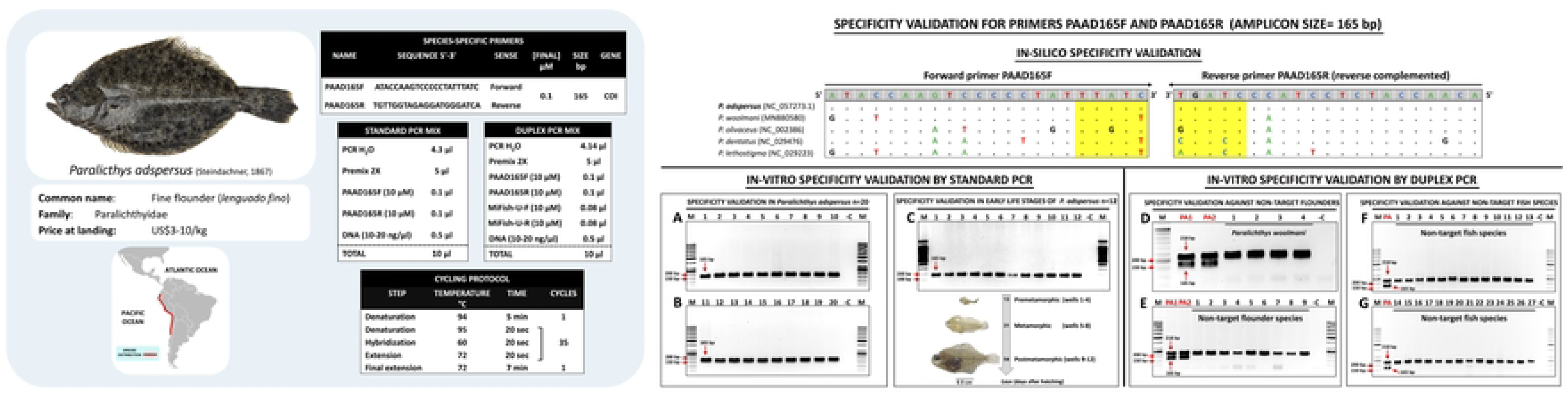
Standardized species-specific primer protocols and validation assay results (in-silico and in-vitro) for the identification of *Paralichthys adspersus*. In the in-silico validation panel, the nucleotides highlighted in yellow correspond to the 3’ ends of each primer hybridization region. Panels A and B: PCR electrophoresis results of the primer set PAAD165F/R showing the positive detection (amplicon size 165 bp) in 20 individuals of *P. adspersus*. Panel C: PCR electrophoresis results of the identification of early life stages of *P. adspersus*. Panels D to G: PCR electrophoresis results of the specificity validation against non-target species by duplex PCR using the primer set PAAD165F/R and an endogenous control amplified by the fish universal primers MiFish-U targeting a partial fragment (218 bp) of the 12S rRNA gene, wells “PA” are positive controls of *P. adspersus*, Panel D: validation against congener *P. woolmani* (wells 1 to 4), E: validation against 5 non-target flatfish species, wells 1-2 *Etropus ectenes*, wells 3-4 *Cyclopsetta querna*, wells 5-6 *Hippoglossina tetrophthalma*, well 7 *Ancylopsetta dendritica*, and wells 8-9 *Symphurus chabanaudi*, panels F and G: validation against non-target fish species listed in S1 Table whose code numbers correspond to the well positions in the agarose gels.

**Fig 6.**
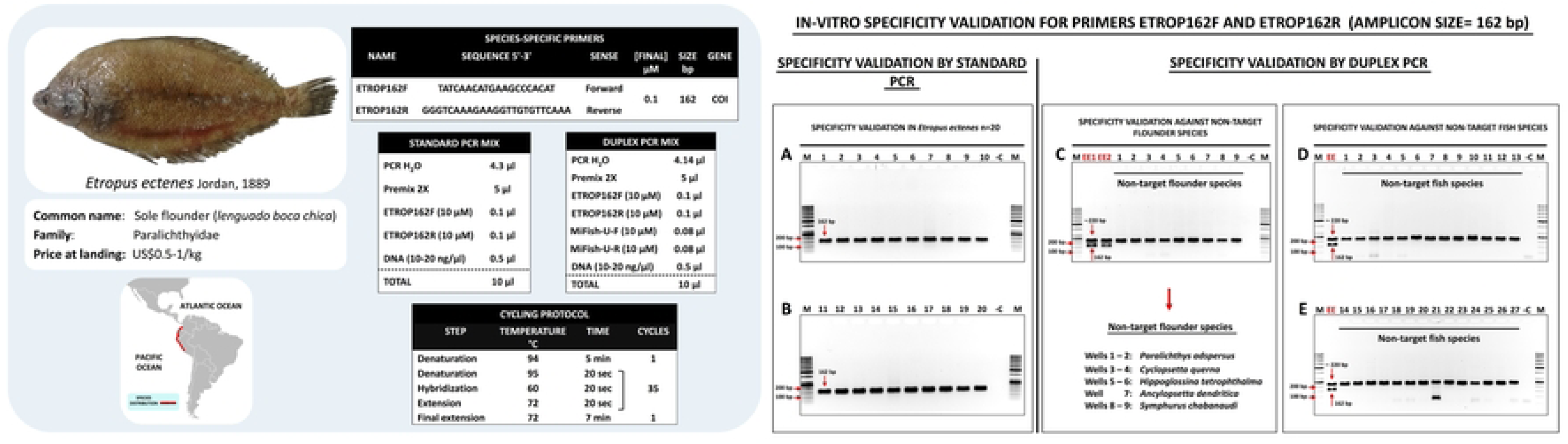
Standardized species-specific primer protocols and in-vitro validation assay results for the identification of *Etropus ectenes.* Panels A and B: PCR electrophoresis results of the primer set ETROP162F/R showing the positive detection (amplicon size 162 bp) in 20 individuals of *E. ectenes*. Panels C to E: PCR electrophoresis results of the specificity validation against non-target species by duplex PCR of the primer set ETROP162F/R and an endogenous control using the fish universal primers MiFish-U targeting a partial fragment (≈ 220 bp) of the 12S rRNA gene where wells “EE” are positive controls of *E. ectenes*, Panel C: validation against 5 non-target flatfish species, Panels D and E: validation against non-target fish species listed in S1 Table whose code numbers correspond to the well positions in the agarose gels.

**Fig 7.**
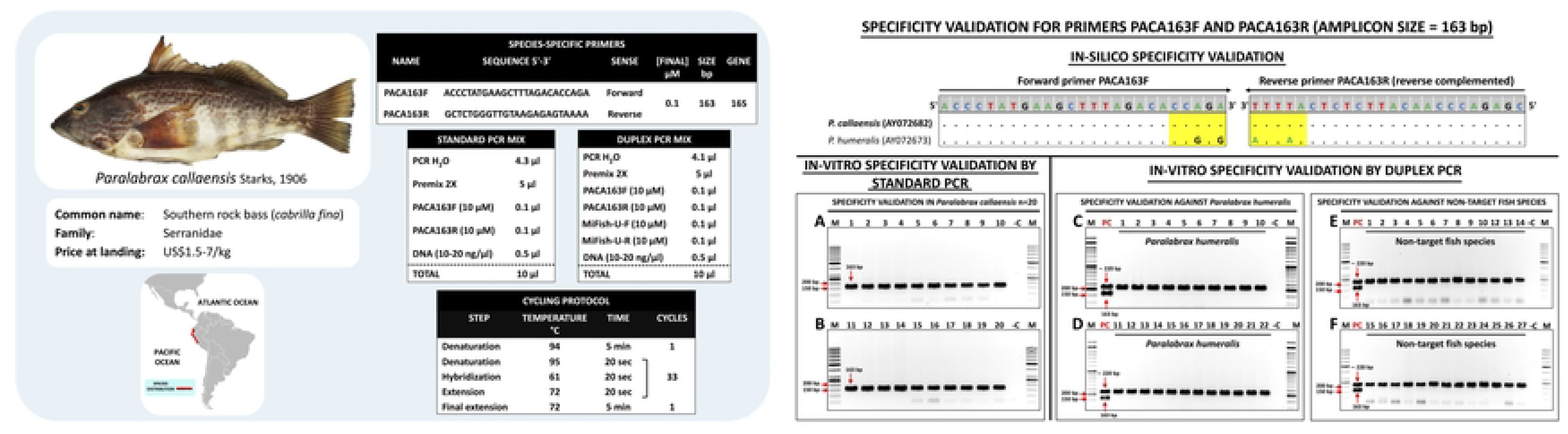
Standardized species-specific primer protocols and validation assay results (in-silico and in-vitro) for the identification of *Paralabrax callaensis*. In the in-silico validation panel, the nucleotides highlighted in yellow correspond to the 3’ ends of each primer hybridization region. Panels A and B: PCR electrophoresis results of the primer set PACA163F/R showing the positive detection (amplicon size 163 bp) in 20 individuals of *P. callaensis*. Panels C to F: PCR electrophoresis results of the specificity validation against non-target species by duplex PCR of the primer set PACA163F/R and an endogenous control using the fish universal primers MiFish-U targeting a partial fragment (≈ 220 bp) of the 12S rRNA gene where wells “PC” are positive controls of *P. callaensis*, panels C and D: validation against 22 individuals of non-target congener *P. humeralis*, panels E and F: validation against non-target fish species listed in S1 Table whose code numbers correspond to the well positions in the agarose gel.

**Fig 8.**
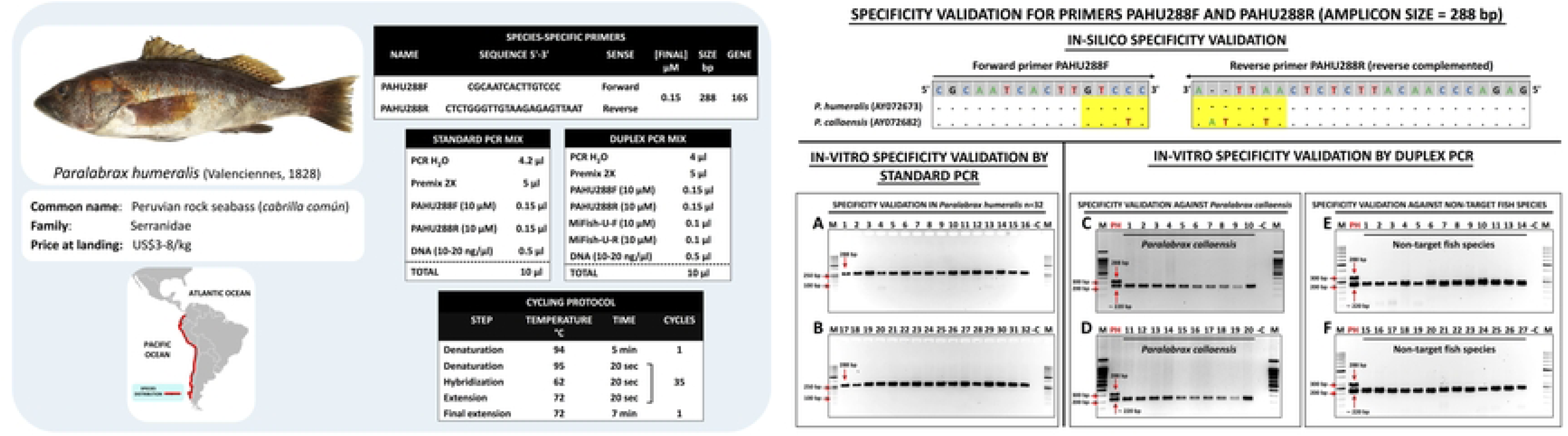
Standardized species-specific primer protocols and validation assay results (in-silico and in-vitro) for the identification of *Paralabrax humeralis*. In the in-silico validation panel, the nucleotides highlighted in yellow correspond to the 3’ ends of each primer hybridization region. Panels A and B: PCR electrophoresis results of the primer set PAHU288F/R showing the positive detection (amplicon size 288 bp) in 32 individuals of *P. humeralis*. Panels C to F: PCR electrophoresis results of the specificity validation against non-target species by duplex PCR of the primer set PAHU288F/R and an endogenous control using the fish universal primers MiFish-U targeting a partial fragment (≈ 220 bp) of the 12S rRNA gene where wells “PH” are positive controls of *P. humeralis*, panels C and D: validation against 20 individuals of non-target congener *P. callaensis*, E and F: validation against non-target fish species listed in S1 Table whose code numbers correspond to the well positions in the agarose gel.

**Fig 9.**
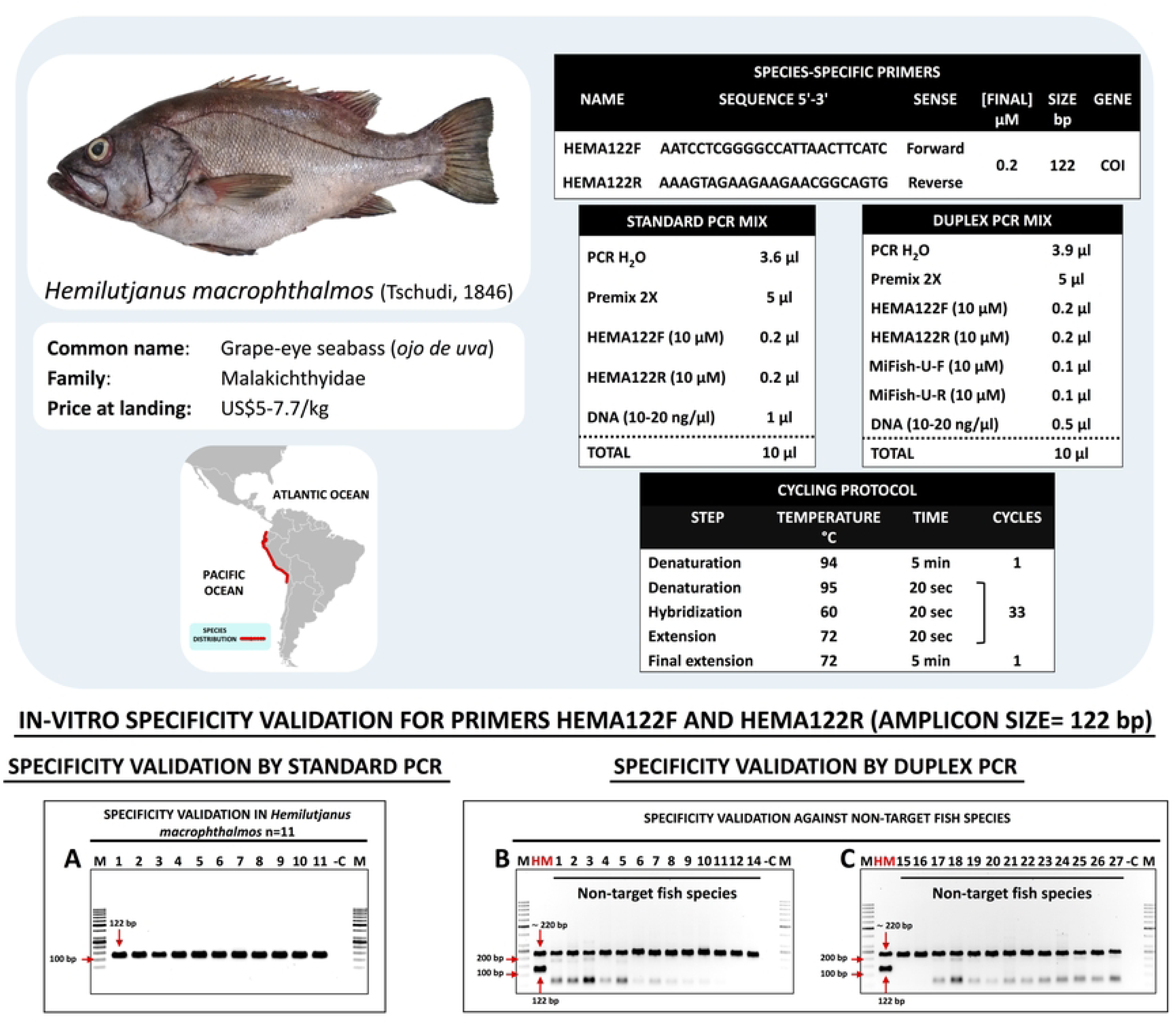
Standardized species-specific primer protocols and in-vitro validation assay results for the identification of *Hemilutjanus macrophthalmos*. Panel A: PCR electrophoresis results of the primer set HEMA122F/R showing the positive detection (amplicon size 122 bp) in 11 individuals of *H. macrophthalmos*. Panels B and C: PCR electrophoresis results of the specificity validation against non-target species by duplex PCR of the primer set HEMA122F/R and an endogenous control using the fish universal primers MiFish-U targeting a partial fragment (≈ 220 bp) of the 12S rRNA gene where wells “HM” are positive controls of *H. macrophthalmos*, non-target fish species are listed in S1 Table whose code numbers correspond to the well positions in the agarose gel.

**Fig 10.**
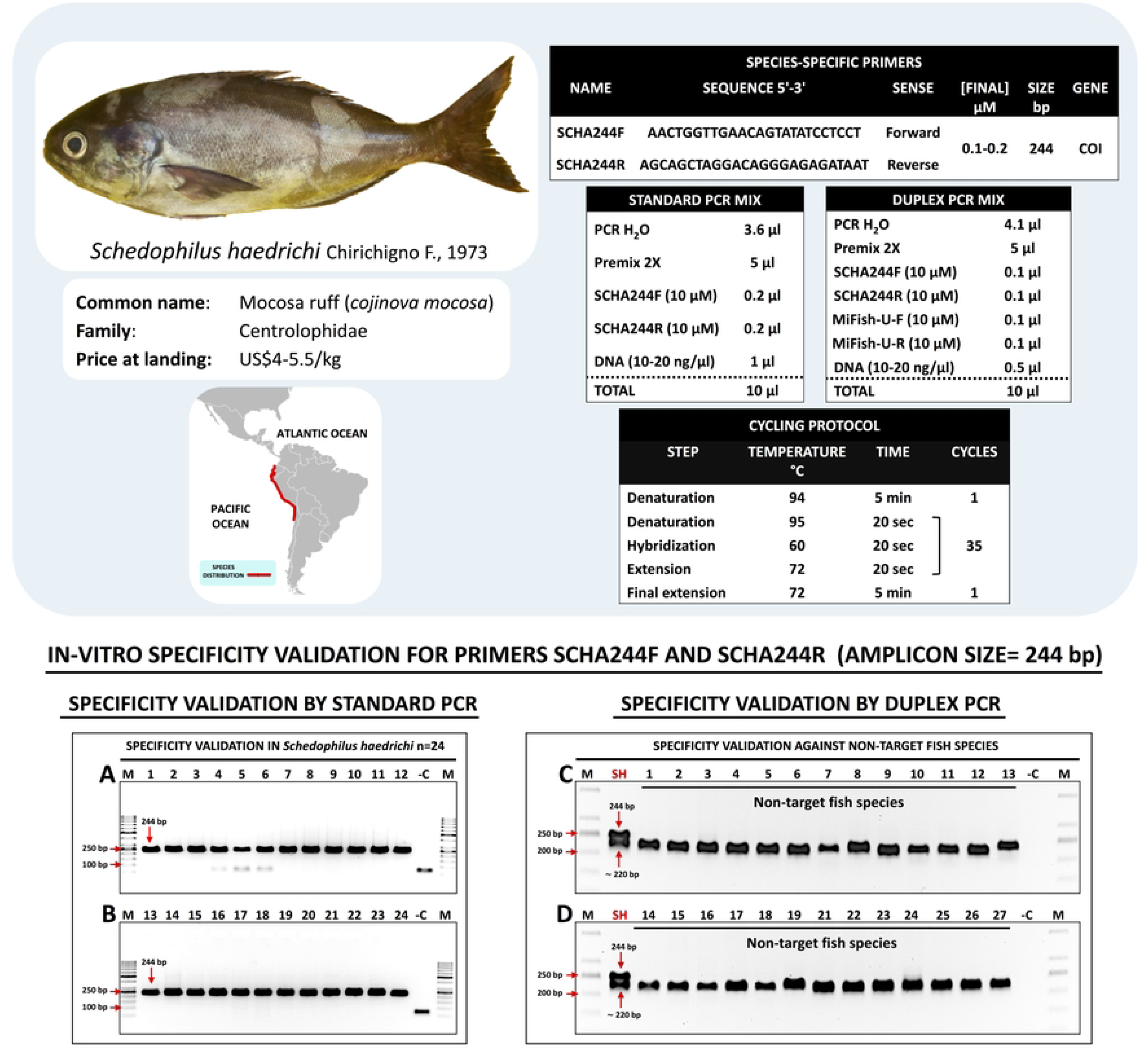
Standardized species-specific primer protocols and in-vitro validation assay results for the identification of *Schedophilus haedrichi*. Panels A and B: PCR electrophoresis results of the primer set SCHA244F/R showing the positive detection (amplicon size 244bp) in 24 individuals of *S*. *haedrichi*. Panels C and D: PCR electrophoresis results of the specificity validation against non-target species by duplex PCR of the primer set SCHA244F/R and an endogenous control using the fish universal primers MiFish-U targeting a partial fragment (≈ 220 bp) of the 12S rRNA gene where wells “SH” are positive controls of *S*. *haedrichi*, non-target fish species are listed in S1 Table whose code numbers correspond to the well positions in the agarose gel.

**Fig 11.**
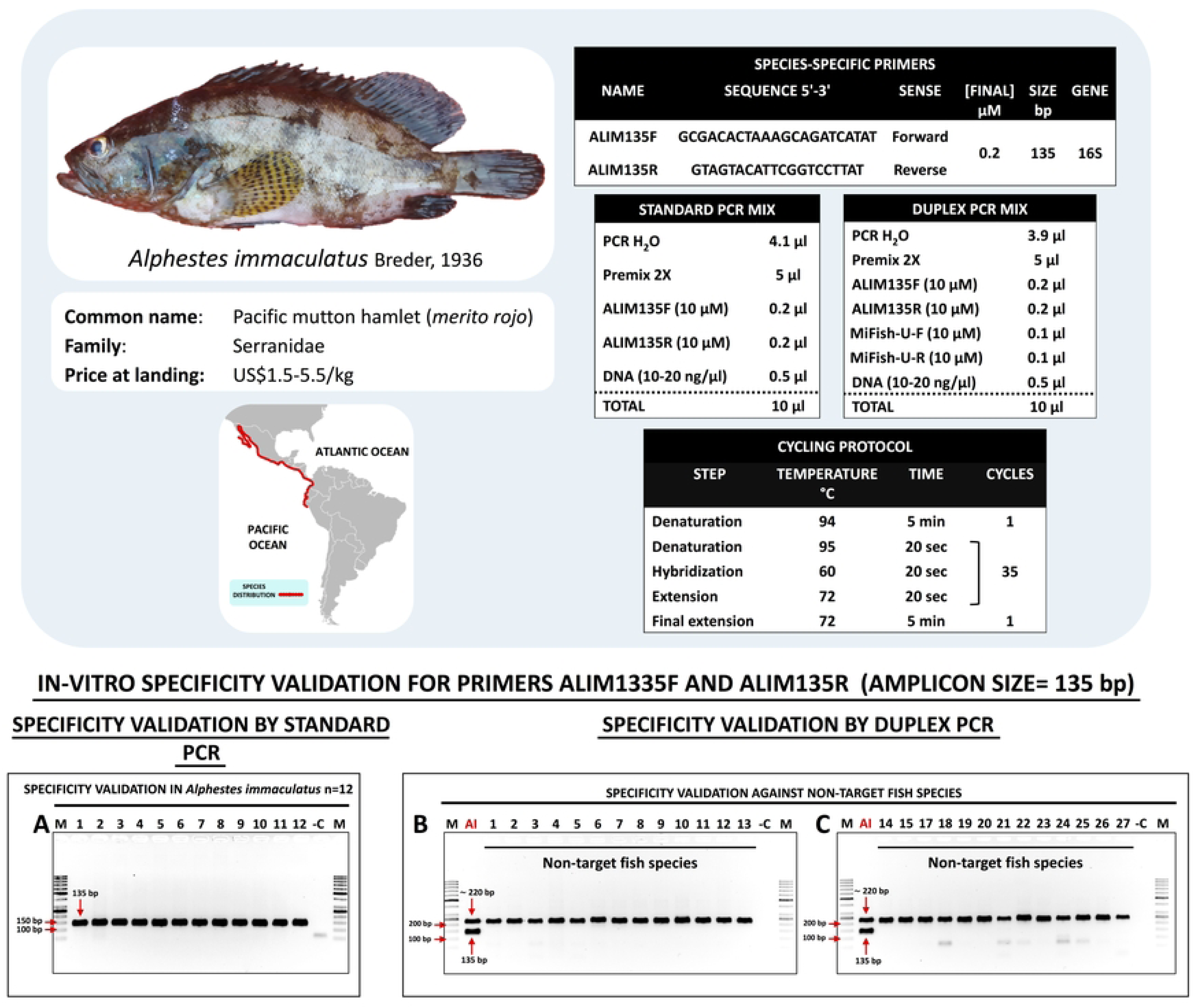
Standardized species-specific primer protocols and in-vitro validation assay results for the identification of *Alphestes immaculatus*. Panel A: PCR electrophoresis results of the primer set ALIM135F/R showing the positive detection (amplicon size 135 bp) in 12 individuals of *A. immaculatus*. Panels B and C: PCR electrophoresis results of the specificity validation against non-target species by duplex PCR of the primer set ALIM135F/R and an endogenous control using the fish universal primers MiFish-U targeting a partial fragment (≈ 220 bp) of the 12S rRNA gene where wells “AI” are positive controls of *A. immaculatus*, non-target fish species are listed in S1 Table whose code numbers correspond to the well positions in the agarose gel.

**Fig 12.**
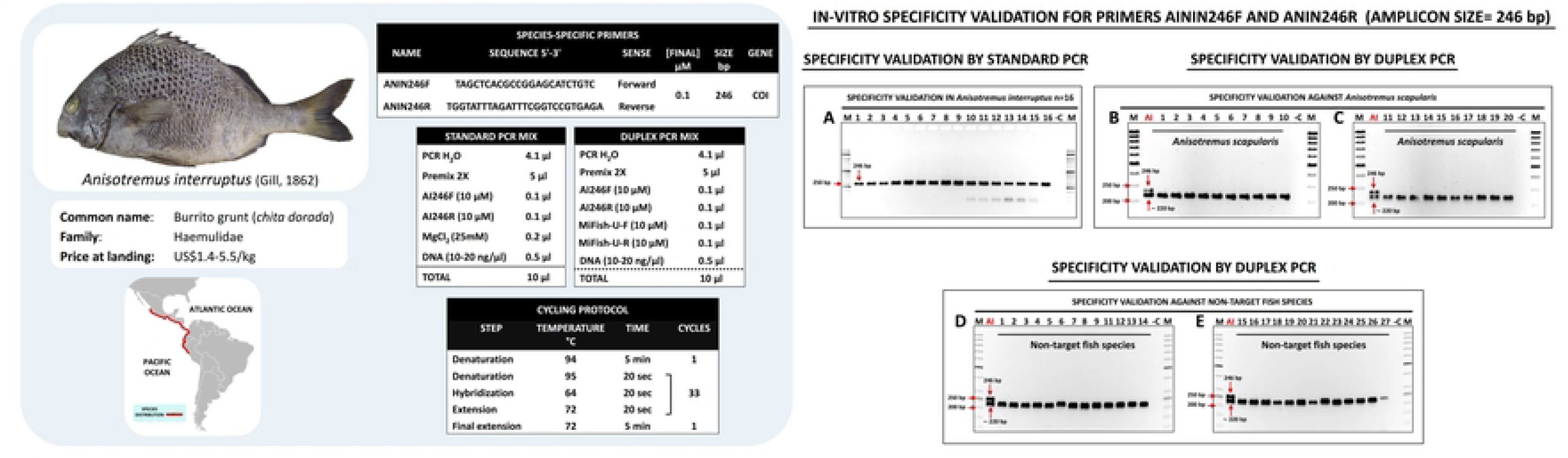
Standardized species-specific primer protocols and in-vitro validation assay results for the identification of *Anisotremus interruptus*. Panel A: PCR electrophoresis results of the primer set ANIN246F/R showing the positive detection (amplicon size 246 bp) in 16 individuals of *A. interruptus*. Panels B to E: PCR electrophoresis results of the specificity validation against non-target species by duplex PCR of the primer set ANIN246F/R and an endogenous control using the fish universal primers MiFish-U targeting a partial fragment (≈ 220 bp) of the 12S rRNA gene where wells “AI” are positive controls of *A. interruptus*, panels B and C: validation against 20 individuals of non-target congener *A. scapularis*, panels D and E: validation against non-target fish species listed in S1 Table whose code numbers correspond to the well positions in the agarose gel.

The SSP set designed for the identification of *A. purpuratus* was evaluated through three stages including in-silico, in-vitro (standard and qPCR using gDNA from target and non-target scallop tissues), and in-situ (qPCR using eDNA from seawater); while the SSPs targeting squid and fish species were validated through in-silico and in-vitro stages (Fig 2). The annealing temperature of the SSPs ranged from 60 to 64 °C, ensuring a high stringency of the PCR and qPCR assays. Targeted PCR products ranged from 129 to 288 bp, which enabled the successful amplification of degraded DNA commonly presented in eDNA samples and processed seafood.

### 3.1 Scallops: *Argopecten purpuratus* (Peruvian scallop)

#### 3.1.1 In-silico SSP validation

The best candidate gene for SSPs design in *A. purpuratus* was found to be the mitochondrial 16S rRNA. To find a region with enough interspecific divergence between closely related *Argopecten* species, we sequenced a partial fragment of the 16S gene in 34 individuals of *A. purpuratus* (GenBank accessions from PP087160 to PP087193) and 27 individuals of its sympatric congener *A. ventricosus* (GenBank accessions from PP087194 to PP087220). After sequence alignment, a common *Argopecten* forward primer “ARGOF” was designed based on a conserved region, whereas a specific *A. purpuratus* reverse primer namely ARPU129R was developed based on the mutations observed in a highly variable region (Fig 3, S2 File). The in-silico specificity validation included all available GenBank sequences from the mitochondrial 16S rRNA gene of *A. purpuratus* (n=34) and sequences determined herein (n=34). The forward primer ARGOF matches perfectly to 67 sequences while only one sequence was found to contain a single mismatch at the twelfth nucleotide from the 5’ end of the primer (S1 Fig, S1 File). The reverse SSP ARPU129R had complete sequence homology with 45 *A. purpuratus* sequences (67%, haplotype “C”), exhibited a single mismatch at the fourth nucleotide from the 3’ end in 21 sequences (30%, haplotype “D”), or contained an additional mismatch (to those found in haplotype “D”) at the fifteenth nucleotide from the 3’ end in two sequences (3%, haplotype “E”) (S1 Fig and S1 File). Individuals containing haplotypes “D” (n=3, Genbank accessions PP087160, PP087169, PP087180) and “E” (n=1, GenBank accession PP087176) were efficiently amplified in further endpoint PCR and qPCR in-vitro analysis. The in-silico specificity validation against non-target species (Fig 3, S2 File) indicated that the forward primer ARGOF contains at least a single mismatch in non-target related Pectinidae species, while the highest number of dissimilarities in binding sites from other scallop species was found in the reverse SSP ARPU129R (from 7 to 11 mismatches and insertions).

#### 3.1.2 In-vitro SSP validation by endpoint PCR and qPCR

The SSP set for *A. purpuratus* (ARGOF/ARPU129R) amplified a single specific PCR product of 129 bp (electrophoresis gels A and B in Fig 3), without primer dimer formation in all tested individuals (n=20) collected from two distant bays located in Piura and Pisco regions. The non-target species PCR validation test resulted in no cross-species reaction in any of the 7 bivalve species that included DNA of a congeneric scallop (*A*. *ventricosus*), two oysters (*Pteria sterna* and *Striostrea prismatica*), a penshell (*Atrina maura*), a clam (*Gari solida*), and two mussels species (*Aulacomya atra* and *Perumytilus purpuratus*) (electrophoresis gel C in Fig 3).

Quantitative PCR results also confirmed the high specificity of the SSP set ARGOF/ARPU129R. A Melting curve analysis of the qPCR products showed a single sharp peak with an average melting temperature (Tm) value of 79.79 °C ± 0.1 in all analyzed *A. purpuratus* samples (panel D in Fig 3), indicating the specific yield of a single qPCR product without primer dimer formation or non-specific products. qPCR specificity validation against non-target bivalve species resulted in no amplification curve formation during the 45 cycles (panel E in Fig 3) confirming the absence of cross-species amplification.

Based on the results of the standard curve analysis (panel F in Fig 3), the LOD of the qPCR for *A. purpuratus* was 3.94 × 10^-4^ ng in 2 μl of DNA template (mean Ct 34.55). A qPCR efficiency of 94.74% was obtained, which is within the acceptable range of highly efficient primers (90–110%) as described in the MIQE Guidelines [57].

#### 3.1.3 Direct detection of *A. purpuratus* by endpoint PCR and qPCR

A schematic representation of the direct qPCR and endpoint PCR detection assay for *A. purpuratus* is depicted in S2 Fig. The swabbing, preservation, and DNA elution steps lasted just 12 min. All fresh and processed samples were successfully identified by our direct qPCR (panel B in S2 Fig) and endpoint PCR (panel C in S2 Fig) assays. The direct qPCR assay displayed positive identification signals between 50-60 min (Ct values ranged from 19.55 to 26.86). On the other hand, the results of direct endpoint PCR assay, which were visualized in agarose gel electrophoresis (panel C in S2 Fig), were obtained in a total time of 150 min.

#### 3.1.4 In-situ SSP qPCR validation, inhibition test, and LOD

The in-situ qPCR validation assay results showed that all positive controls (tissue-derived DNA, run in triplicate) included in each qPCR run were successfully amplified with Ct values corresponding to its DNA concentration, whereas the negative control replicates (qPCR water as DNA template and filtered distilled water from eDNA field surveys) gave no fluorescence signal, validating the qPCR assays. Validation of the positive qPCR products using Sanger sequencing confirmed that all eDNA amplicons belonged to *A. purpuratus* (GenBank accessions from PP087144 to PP087154, 100% identity match with *A. purpuratus* mitogenome reference sequence KF601246). Environmental DNA of the Peruvian scallop was detected in all replicates (3/3) from all sampling stations of Sechura Bay (map B in Fig 1), Samanco Bay (map C in Fig 1), and Tortugas Bay (map D in Fig 1), except for La Cruz offshore station in Tumbes region (map A in Fig 1), which is out of the natural distribution range of *A. purpuratus* (Peña, 2001). eDNA concentrations ranged from 2.69×10^-1^ ng (Parachique station) to 8.52×10^-4^ ng (Tortugas 5 station). Parachique station (Sechura Bay) was the only location from where eDNA samples were collected at three different layers of the water column (bottom, midwater, and surface) and where the presence of the Peruvian scallop was confirmed by visual inspection during bottom water sampling. *A. purpuratus* eDNA concentrations from that station were similar among bottom (1.68×10^-2^ ng, Ct 26.53 ± 0.11), middle (2.69×10^-1^ ng, Ct 24.3 ± 0.03), and surface (2.0×10^-1^ ng, Ct 24.75 ± 0.06) levels. eDNA extractions from Tortugas 1 and Tortugas 3 obtained from half filters generated lower *A. purpuratus* eDNA amounts than those extractions using quarter filters (Table 3), possibly due to the presence of inhibitors in those samples.

**Table 3.** eDNA identification results of the species-specific primers targeting *A. purpuratus*. Information from each sampling locations, collection dates and season, geographic coordinates, filtered water volume and level of the water column (B: bottom, M: midwater, S: surface), and qPCR results. Sampling stations with an asterisk (*) denotes eDNA samples extracted from a quarter filter

| Location | Site information | Sampling station | Geographic coordinates | Sampling date (Season) | Vol. (L) | Level | *A. purpuratus* (GenBank accession) | qPCR Ct | Concentration (ng) |
| --- | --- | --- | --- | --- | --- | --- | --- | --- | --- |
| Sechura Bay (Piura region) | Open bay with wild populations of *A. purpuratus* and *A. ventricosus*. This bay is also used for *A. purpuratus* farming (mainly sea ranching, hanging lantern nets to a lesser extent) | Parachique | 05°45'9.7"S, 080°53'52.6"W | October 14, 2017 (Spring) | 0.5 | B | + (PP087144) | 26.53 ± 0.11 | 1.68×10<sup>-2</sup> ± 1.13×10<sup>-3</sup> |
|  |  |  |  |  | 0.5 | M | + (PP087145) | 24.3 ± 0.03 | 2.69×10<sup>-1</sup> ± 5.55×10<sup>-3</sup> |
|  |  |  |  |  | 0.5 | S | + (PP087146) | 24.75 ± 0.06 | 2.0×10<sup>-1</sup> ± 7.36×10<sup>-3</sup> |
|  |  | Bayóvar | 05°48'25.8"S, 081°00'34.9"W | May 11, 2022 (Autumn) | 1 | B | + (PP087147) | 28.42 ± 0.11 | 2.13×10<sup>-2</sup> ± 1.47×10<sup>-3</sup> |
|  |  | Matacaballo | 5°38'10.2"S 80°51'40.7"W | May 12, 2022 (Autumn) | 1 |  | + (PP087148) | 32.67 ± 0.08 | 1.60×10<sup>-3</sup> ± 7.99×10<sup>-5</sup> |
| Samanco Bay (Ancash region) | Semi-enclosed bay with wild and cultured populations (hanging lantern nets) of *A. purpuratus* | El Dorado | 09°12'24"S, 078°31'46"W | February 12, 2019 (Summer) | 1 | S | + (PP087149) | 29.26 ± 0.21 | 6.41×10<sup>-3</sup> ± 8.19×10<sup>-4</sup> |
| Tortugas Bay (Ancash region) | Semi-enclosed bay with wild populations of *A. purpuratus* | Tortugas 1 | 09°22'06.8"S, 078°25'03.3"W | April 29, 2022 (Autumn) | 1 | B | + (PP087150) | 32.14 ± 0.14 | 1.36×10<sup>-3</sup> ± 1.13×10<sup>-4</sup> |
|  |  | Tortugas 1* |  |  |  |  | + (PP087151) | 27.75 ± 0.08 | 7.74×10<sup>-3</sup> ± 3.89×10<sup>-4</sup> |
|  |  | Tortugas 2 | 09°21'38.2"S, 078°25'00.6"W |  | 1 |  | + (PP087152) | 32.94 ± 0.26 | 8.52×10<sup>-4</sup> ± 1.36×10<sup>-4</sup> |
|  |  | Tortugas 3 | 09°21'42.5"S, 078°25'10.4"W |  | 1 |  | + (PP087153) | 31.28 ± 0.24 | 1.86×10<sup>-3</sup> ± 2.60×10<sup>-4</sup> |
|  |  | Tortugas 3* |  |  |  |  | + (PP087154) | 29.34 ± 0.22 | 2.88×10<sup>-3</sup> ± 3.75×10<sup>-4</sup> |
| La Cruz (Tumbes Region) | Offshore sampling station without historical records of *Argopecten purpuratus* | La Cruz | 3°34'35.1"S, 80°37'3.3"W | December 12, 2017 (Spring) | 1 | S | - | - | - |

The eDNA inhibition test indicated that all blank control samples obtained in all sampling stations and the eDNA samples collected at Parachique station (Sechura Bay) showed a Ct shift < 2 compared to that of the positive exogenous DNA control (*A. gigas*), suggesting the absence of inhibitory substances. All the remaining eDNA samples displayed a Ct shift > 2 and therefore were deemed inhibited. In an attempt to overcome this issue, inhibited samples were diluted (1:1, 1:5, 1:10, and 1:20) and qPCR amplified using 0.5 μl of bovine serum albumin (BSA, 20 mg/ml), resulting in positive detections of the target species in all inhibited samples. The LOD of our in-situ eDNA assay for *A. purpuratus* was 2×10^-5^ ng in 2 μl of eDNA template (Ct 35.62), with a qPCR efficiency of 95.87% (Panel I in Fig 3).

### 3.2 Squids: *Doryteuthis gahi* (Patagonian longfin squid)

#### 3.2.1 In-silico and in-vitro SSP validation by endpoint PCR

The SSP set DOGA136F/R amplified a partial fragment of 136 bp within the mitochondrial COI gene of *D. hagi*. We deliberately introduced a “GG tail” on the 5’ ends of both forward and reverse primers to meet the minimum percentage of recommended GC content (40%) to ensure a more stable primer/template binding [63] during PCR amplifications. The specificity of the SSP set DOGA136F/R was supported by the in-silico evaluation using 26 available sequences of the target species (see S3 File) showing perfect complementarity with template strands (disregarding the introduced “GG tail”), while only one individual had a single mismatch in the fifth position from the 3’ end of the forward primer binding site. The in-silico analysis against non-target squid species (Fig 4) detected the presence of 1 to 4 mismatches in the 3’ end of the forward primer in all 8 non-target squid species. A further in-silico test against 142 sequences of *D*. *gigas* available in GenBank/BOLD databases (S4 File) indicated that the potential of false positive yield in *D*. *gigas* DNA is unlikely due to the presence of mismatches in the 3’ end of forward and reverse primers’ hybridization sites. The Primer-BLAST results indicated that *D. hagi* is the best match for the SSP set DOGA136F/R. The same analysis also indicated that the SSPs DOGA136F/R would amplify fragments of 794 and 2546 bp in the diamond squid *Thysanoteuthis rhombus* (Thysanoteuthidae). However, this elusive cosmopolitan squid species is only targeted in Japanese waters [64], while in Peru has no commercial value with few reported bycatch incidents [65].

The in-vitro specificity validation results demonstrated the high performance of the SSP set DOGA136F/R, electrophoresis gel A in Fig 4 shows that 100% of tested target individuals (n=14) were positive, while electrophoresis gel B in Fig 4 demonstrated that no cross-species reactions occurred when tested against 4 non-target squid species including two economically important species from Peru: *L. diomedeae* and *D. gigas*.

### 3.3 Fish: Alphestes immaculatus, Anisotremus interruptus, Etropus ectenes, Hemilutjanus macrophthalmos, Paralabrax callaensis, P. humeralis, Paralichthys adspersus, and Schedophilus haedrichi

#### 3.3.1 In-silico and in-vitro SSP validation by endpoint PCR

Based in the in-silico analysis of mitochondrial genes for SSP design for fish species, the COI gene was the best candidate in *P. adspersus* (Fig 5, S5 File), *E. ectenes* (panel A in S4 Fig), *H. macrophthalmos* (panel B in S4 Fig), *S. haedrichi* (panel C in S4 Fig), and *A. interruptus* (panel E in S4 Fig), while the 16S rRNA gene was more suitable in *P. callaensis* (Fig 7), *P. humeralis* (Fig 8), and *A. immaculatus* (panel D in S4 Fig).

The intraspecific specificity of all putative SSPs for the 8 selected fish species was evaluated using self-generated and reference sequences available in GenBank database, which ranged from 4 sequences in *E. ectenes* to 175 sequences in *A. interruptus*. The SSP sets for *P. callaensis* (PACA163F/R, S8 File), *P. humeralis* (PAHU288F/R, S9 File), *H. macrophthalmos* (HEMA122F/R S10 File), *S. haedrichi* (SCHA244F/R, S11 File), and *A. immaculatus* (ALIM135F/R, S12 File) showed a perfect match (100% complementarity) in all analyzed reference sequences from each target species, disregarding the two last nucleotides of the 5’ end of the forward primer ALIM135F where we deliberately introduced a “GC tail” to meet the minimum percentage of recommended GC content (40%) ensuring a more stable primer/template binding [63]. On the other hand, a single mismatch mutation was detected in one primer targeting the following species: *P. adspersus* (PAAD165R, a mismatch at the fourth nt position from the primer’s 3’ end was found in 2 out of 24 analyzed sequences, S5 File), *E. ectenes* (ETROP162R, a mismatch at the fourth nt position from the primer’s 3’ end, S7 File), and *A. interruptus* (ANIN246F, a mismatch at the 10 nt position of the primer’s 3’ end, S13 File).

All in-silico validation assays against not-target species showed the presence from one to several mismatches positioned along the primer hybridization regions including the 3’ end, which is known to result in the most detrimental effects for polymerase activity [11, 12]. In-silico results for *P. adspersus* (PAAD165F/R) are shown in Fig 5 and S6 File, for *E. ectenes* (ETROP162F/R) in panel A of S4 Fig, for *H. macrophthalmos* (HEMA122F/R) in panel B of S4 Fig, for *S. haedrichi* (SCHA244F/R) in panel C of S4 Fig, for *A. immaculatus* (ALIM135F/R) in panel D of S4 Fig, for *A. interruptus* (ANIN246F) in panel E of S4 Fig, for *P. callaensis* (PACA163F/R) in Fig 7, and for *P. humeralis* (PAHU288F/R) in Fig 8.

The Primer-BLAST search indicated that the SSP sets designed for *A. interruptus*, *A. immaculatus*, *H. macrophthalmos*, *P. adspersus*, *P. humeralis*, and *S. haedrichi* showed the best match result only with sequences from their target species. The primer set PAHU288F/R targeting *P. humeralis* showed a single mismatch in the hybridization region of the reverse primer of its congeneric relative *P. nebulifer*, which occurs in Northern Eastern Pacific from Santa Cruz in central California (USA) to Magdalena Bay in Baja California (Mexico) [66]. The primer set ETROP162F/R was originally designed based on the mitochondrial COI sequence of *E. crossotus*, due to the lack of reference sequences of *E. ectenes* at the moment of primer design. Both species co-occur in northern Peru [41], however only *E. ectenes* is reported to species level in official landing records [67–69].

The in-vitro specificity validation assays were evaluated using 11 to 32 specimens of the target fish species, resulting in the specific amplification of a single specific amplicon in all tested individuals. Thus, species-specific PCR products of the following sizes were obtained for each target species: 165 bp in 32 specimens of *P. adspersus* corresponding to 20 adults and 12 larvae/juveniles (electrophoresis gels A to C in Fig 5), 162 bp in 20 specimens of *E. ectenes* (electrophoresis gels A and B in Fig 6), 163 bp in 20 specimens of *P. callaensis* (electrophoresis gels A and B in Fig 7), 288 bp in 32 specimens of *P. humeralis* (electrophoresis gels A and B in Fig 8), 122 bp in 11 specimens of *H. macrophthalmos* (electrophoresis gel A in Fig 9), 244 bp in 24 specimens of *S. haedrichi* (electrophoresis gels A and B in Fig 10), 135 bp in 12 specimens of *A. immaculatus* (electrophoresis gel A in Fig 11), and 246 bp in 16 specimens of *A. interruptus* (electrophoresis gels A in Fig 12). Finally, the results from the duplex PCR in-vitro specificity assays against non-target fish species demonstrated the high specificity of all the SSPs developed herein. All non-target fish species (listed in Table S1) amplified only the endogenous control band of about 220 bp belonging to the 12S rRNA gene, while only the positive control samples (DNA from target species) yielded two bands corresponding to the species-specific amplicon and the endogenous control band (electrophoresis gels are shown in Fig 5 to Fig 12). Furthermore, specificity assays against non-target congeneric species were evaluated using the SSPs targeting *Paralichthys adspersus*, *Paralabrax callaensis*, *Paralabrax humeralis*, and *Anisotremus interruptus* using DNA from congeneric *Paralichthys woolmani* (n=4, electrophoresis gel D in Fig 5), *P. humeralis* (n=22, electrophoresis gels C and D in Fig 7), *P. callaensis* (n=20, electrophoresis gels C and D in Fig 8), and *Anisotremus scapularis* (n=20, electrophoresis gels B and C in Fig 12), respectively. No cross-reactions were observed at any of the DNA samples belonging to congeneric species.

### 3.9 Application of SSP to authenticate commercial cooked fish samples from restaurants

Our SSP assays allowed us to authenticate target species in cooked samples containing *A. purpuratus* (seafood rice, n=1), *D. gahi* (seafood rice n=2, and cebiche n=1), and *Anisotremus interruptus* (fried n=1) (Table 4). Endpoint PCR conditions for the authentication of dishes containing scallops, squids, and grunts were the same as shown in Fig 3, Fig 4, and Fig 12 respectively. Four additional fish species were identified in two independent multiplex PCR assays. A first multiplex PCR (Panel A in S3 Fig), which included SSPs for the two congeneric species *P. callaensis* (PACA163F/R) and *P. humeralis* (PAHU288F/R), enabled the identification of four samples (cebiche n=1, fried n=1, stewed n=2) based on the species-specific size band (163 bp for *P. callaensis* and 288 bp for *P. humeralis*). Species identities of PCR products from cooked “*cabrilla*” samples were further verified by Sanger sequencing (GenBank accessions PP087227 to PP087229). A second multiplex assay (panel B in S3 Fig) containing SSPs for two species (*H. macrophthalmos* HEMA122F/R and *S. haedrichi* SCHA244F/R) commonly marketed as “*ojo de uva*” (Spanish for grape-eye) enabled the simultaneous identification of DNA from both species in a stewed *H. macrophthalmos* sample (122 bp) and a *S. haedrichi* ceviche sample (244 bp).

**Table 4.** Authentication of commercial cooked seafood samples collected from Peruvian restaurants.

Species-specific primers for the identification of economically important marine resources from Peru
| Sample number | Sample label<br>Spanish<br>(English) | Date | Presenta<br>tion | Locality<br>(Region) | Species-Specific<br>Primer set | Ampli<br>con<br>size<br>(bp) | Rapid molecular<br>diagnosis |
| --- | --- | --- | --- | --- | --- | --- | --- |
| 1 | Cabrilla<br>(rock<br>seabass) | May 7,<br>2019 | Fried | Piura (Piura) | PACA163F/R* +<br>PAHU288F/R | 163 | *Paralabrax*<br>*callaensis* |
| 2 | Cabrillon<br>(jumbo<br>rock<br>seabass) | May 7,<br>2019 | Stewed | Piura (Piura) | PACA163F/R* +<br>PAHU288F/R | 163 | *Paralabrax*<br>*callaensis* |
| 3 | Cabrilla<br>(rock<br>seabass) | May 7,<br>2019 | Cebiche | Chimbote<br>(Ancash) | PACA163F/R +<br>PAHU288F/R* | 288 | *Paralabrax*<br>*humeralis* |
| 4 | Cabrilla<br>(rock<br>seabass) | May 7,<br>2019 | Stewed | Chimbote<br>(Ancash) | PACA163F/R +<br>PAHU288F/R* | 288 | *Paralabrax*<br>*humeralis* |
| 5 | Calamar<br>(squid) | May 7,<br>2019 | Seafood<br>rice | Piura (Piura) | DOGA136F/R | 136 | *Doryteuthis gahi* |
| 6 | Berrugata | May 10,<br>2019 | Fried | Piura (Piura) | ANIN246F/R | 246 | *Anisotremus*<br>*interruptus* |
| 7 | Ojo de uva<br>(grape-eye<br>seabass) | May 11,<br>2019 | Cebiche | Huanchaco<br>(La Libertad) | SCHA244F/R* +<br>HEMA122F/R | 244 | *Schedophilus*<br>*haedrichi* |
| 8 | Concha de<br>abanico<br>(Peruvian<br>scallop) | March<br>24, 2024 | Seafood<br>rice | Chimbote<br>(Ancash) | ARGOF +<br>ARPU129R | 129 | *Argopecten*<br>*purpuratus* |
| 9 | Ojo de uva<br>(grape-eye<br>seabass) | April 22,<br>2024 | Stewed | Tumbes<br>(Tumbes) | SCHA244F/R +<br>HEMA122F/R* | 122 | *Hemilutjanus*<br>*macrophthalmos* |
| 10 | Calamar<br>(squid) | April 24,<br>2024 | Cebiche | Chimbote<br>(Ancash) | DOGA136F/R | 136 | *Doryteuthis gahi* |
| 11 | Calamar<br>(squid) | April 24,<br>2024 | Seafood<br>rice | Chimbote<br>(Ancash) | DOGA136F/R | 136 | *Doryteuthis gahi* |

## 4. Discussion

This study developed and validated different SSP detection assays targeting 10 commercially important fish and shellfish species from Peru and evaluated them in different scenarios including eDNA monitoring, early-life stage identification, and authentication of seafood products. The novelty of this study not only lies in the wide variety of starting genetic material analyzed during the performance and utility evaluations of our novel SSPs (DNA from early life stages and adult individuals, forensic samples, commercial cooked and processed products, environmental DNA from seawater, and tissue swabbing) but also in the different PCR approaches that were used for the molecular detections (endpoint PCR, multiplex PCR, qPCR, direct PCR and qPCR, eDNA qPCR, and DNA sequencing) (Fig 2).

The Peruvian scallop *A*. *purpuratus* represents the most important scallop species along the Pacific coast of South America [70, 71]. Its distribution ranged from Paita in Peru (5° S) to Valparaíso in Chile (33° S) [72] where is commercially exploited, with Peruvian production shipped to at least 19 countries [73]. The SSP assay targeting *A. purpuratus* was validated via endpoint PCR and qPCR methods, resulting in high specificity and efficiency in amplifying tissue-extracted DNA from fresh and processed commercial samples and swab samples without the need for DNA isolation (direct PCR, S2 Fig). Moreover, no cross-species reactions were observed during the PCR and qPCR in-vitro validations against non-target bivalve species, thereby demonstrating its potential applicability to detect mislabeling. The Peruvian scallop has been documented to be involved in substitution and mislabeling in different countries. For example, Parrondo et al. [74] identified cooked and frozen *A. purpuratus* samples incorrectly labeled as “*zamburiña*” (Spanish vernacular name for *Mimachlamys varia*) or “*vieira*” (*Pecten maximus*) in Spanish restaurants and supermarkets. Similarly, Näumann et al. [75] and Klapper and Schröder [8] detected *A. purpuratus* mislabeled as *Pecten* spp. in commercial samples from Germany. The in-silico test against 20 commercial non-target scallop species (S2 File) predicted that cross-species reaction of our SSPs is unlikely, even with its phylogenetically closest living relative *A. ventricosus* [76, 77], which was confirmed by the in-vitro analysis where no cross-amplification was observed in DNA samples of *A. ventricosus* (electrophoresis gel C in Fig 3). Therefore, we expect that the SSPs for *A. purpuratus* will not cross-react with other more distant related scallop species not included in our in-vitro assays. However, further analysis using more scallop species’ DNA will be needed to confirm the in-silico results obtained herein.

The high specificity of the Peruvian scallop’s SSPs was further demonstrated in a novel eDNA assay, which represents the first effort to evaluate the potential of single-species eDNA-based biomonitoring technology in Peruvian marine ecosystems. The scallop eDNA assay successfully detected the presence of *A. purpuratus* in all eDNA sampling sites where it is known to occur (maps B, C, and D in Fig 1). During the collection of bottom eDNA samples from Parachique station (Sechura Bay), we spotted wild stocks of *A. purpuratus* and *A. ventricosus* occurring in sympatry. Nevertheless, our eDNA assay only detected the target scallop species, without signals of cross-reaction with eDNA of its congeneric relative *A. ventricosus*. This result emphasizes the high specificity of our assay and therefore it can be employed to discover new Peruvian scallop beds or to get further insights into population dynamics during extreme atmospheric conditions such as ENSO, which is known to cause notorious changes in the production and dispersal of the Peruvian scallop larvae [78].

Due to its benthic nature, we expected to find higher *A. purpuratus* eDNA concentrations in bottom samples. However, estimated eDNA amounts from bottom, midwater, and surface samples from Parachique station were similar (Table 3). A possible reason for that may be that all eDNA samples from Parachique station were collected in shallow waters (from 0 to 6 m depth), therefore it is likely that there was little vertical stratification. A plausible explanation for the slightly higher eDNA concentrations observed in midwater and surface samples would be that planktonic gamete/larvae were present in the water column above the bottom layer presumably as a result of spawning activity occurring the days before and/or during the collection of water samples. eDNA samples from Parachique station (Sechura Bay) were collected in October 2017 which corresponds to the peak spring spawning period (September to November) reported for Sechura Bay’s Peruvian scallop populations [79]. However, we cannot rule out the possibility that restocking activity in nearby bottom culture sites may have also accounted for the eDNA levels observed in our results, due to the natural horizontal dispersion process of the eDNA [80] shed by the newly introduced spat. A strong Coastal El Niño event took place from January to April 2017 [70, 81] causing high mortality rates in *A. purpuratus* populations from Sechura Bay. Aiming to overcome the significant economic loss caused by the massive mortality event, some scallop producers may have used wild or hatchery-produced spat to restock their farms. Further studies based on repeated temporal sampling from wild scallop population sites (without aquaculture activity) and supported by quantitative assessment of gamete production and mesocosm experiments are needed to confirm the utility of our eDNA assay to identify space-temporal spawning events in Peruvian scallop populations.

The Patagonian longfin squid *D. gahi* is targeted by artisanal fisheries in Argentina, Chile, and Peru [82] and exported to several countries around the world [73, 83]. In several nations, *D. hagi* (Loliginidae) and *Dosidicus gigas* (Ommastrephidae) are frequently marketed using the generic term “squid” or “*calamar*” (in Spanish-speaking countries). However, *D. gigas* is of lower economic value with year-round availability which makes it the best candidate to substitute more expensive Loliginidae squids [84, 85]. The specificity of the SSP set DOGA136F/R targeting *D. hagi* was supported by the in-silico (S3 File and S4 File) and in-vitro specificity analysis performed in *D. hagi* samples (electrophoresis gel A in Fig 4, Table 4) and against 4 non-target squid species of commercial value namely *Doryteuthis opalescens*, *Lolliguncula diomedeae*, *Dosidicus gigas*, and *Todarodes pacificus* (electrophoresis gel B in Fig 4). Considering all these results together, the squid detection assay validated herein will offer a rapid and economical method to authenticate *D. hagi* products.

In Peru, several species of the family Paralichtyidae are simply sold as “*lenguado*” (Spanish for flounder) [38, 41]. Among them, the fine flounder (*P. adspersus*) and the sole flounder (*E. ectenes*) represent two important Peruvian flatfish species of commercial interest. The former species not only supports the most important Peruvian artisanal flatfish fishery but is also the most demanded and expensive flounder species [86] and known to be a target of mislabeling [43, 46]. The in-vitro PCR test against 32 non-target species that included 6 commercially important flatfish species (electrophoresis gels D to G in Fig 5) demonstrated that our primer set PAAD165F/R is specific for *P. adspersus*. In-vitro results of the primer set ETROP162F/R showed that it only reacted with DNA from *E. ectenes* (electrophoresis gels C to E in Fig 6), although it is also expected to hybridize with DNA from its congener *E. crossotus*, as mentioned in Results section. The in-silico validation against 10 congeneric *Paralichthys* species of global commercial value indicated that cross-species reaction is unlikely due to the presence of mismatches in the 3’ end of both primers in non-target species (S6 File), suggesting the potential utility of our assay in international markets. However, further in-vitro analysis including gDNA of other *Paralichthys* species will be required to confirm our in-silico findings.

Our results also demonstrated that the SSPs for *P. adspersus* can accurately identify different early life stages (electrophoresis gel C in Fig 5). Further analyses using unsorted plankton samples are required to evaluate the performance of the SSP set PAAD165F/R in detecting early life stages of the target species in the presence of background plankton community. Molecular assays for the accurate identification of early life stages of commercial fish species can provide support in spawning ground detection and seasonal changes in spawning activity [87], and during wild egg/fry collection for farming purposes limiting accidental rearing of non-target species [88].

Designing efficient SSPs to discriminate closely related species can be challenging, especially when sequence differences in the primer hybridization region are limited to 1 or 2 positions. Purine-purine (i.e., A-A, A-G, G-G, or G-A) and pyrimidine-pyrimidine (i.e., C-C) mismatch pairings are known as critical mismatches that have the most detrimental effects when positioned at the primer’s 3’ terminal [11, 12]. In addition to positioning mismatches within the primer’s 3’ end region, other variables can be adjusted to bias amplifications in favor of the target sequence including lower primer concentrations, fewer PCR cycles, and raising the annealing temperature as much as possible [89, 90]. Furthermore, it is highly suggested the use of commercial PCR kits containing ammonium sulfate on its buffer system, since it enhances the primer specificity, decreasing the chances of primer dimer formation and the yield of non-specific PCR products [56]. Herein, we demonstrated that the punctual mutations and indels present at the 3’ end of the primer’s hybridization region of non-target congeneric species *P. humeralis* (Fig 7) and *P. callaensis* (Fig 8) combined with high annealing temperatures (61 and 62 °C respectively) and low primer concentrations (0.1 and 0.15 μM respectively) were proven to be effective in hampering cross-species reactions. The two *Paralabrax* species targeted in our study support important artisanal fisheries in Peru [91] where they are frequently marketed as “*cabrilla*” [38] (authors’ personal observation). We were able to standardize a multiplex PCR assay combining the discriminatory power of both SSP sets PACA163F/R (targeting *P. callaensis*) and PAHU288F/R (*P. humeralis*) in a single reaction, with which we could rapidly and accurately identify cooked samples belonging to both target species, as shown in panel A of S3 Fig.

We successfully developed a second multiplex PCR that was proven effective in the simultaneous identification of cooked presentations of two non-related fish species that share the same market name “*ojo de uva*” (grape-eye) namely *H. macrophthalmos* (Malakichthyidae) and *S. haedrichi* (Centrolophidae) (panel B in S3 Fig). The latter species is also known as “*mocosa*”, “*cojinoba del norte*”, and “*cojinoba mocosa*” [41, 69]. However, in some regions from northern Peru (e.g. La Libertad and Lambayeque regions) the commercial name “*ojo de uva*” is also attributed to *S. haedrichi* (authors’ personal observations), which does not necessarily imply mislabeling or species substitution, but rather a case of synonymy with the common name of *H*. *macrophthalmos*. In this regard, our multiplex assay targeting both species marketed as “*ojo de uva*” can be effectively applied for their simultaneous differentiation (panel B in S3 Fig). In Peru, *H. macrophthalmos* is considered a luxury commodity with tight supply and in high demand, especially by high-end restaurants. High historic catches of this species were reported in 1987 and 1996, but it was drastically lower after 2014 suggesting overexploitation [44, 92]. To ensure the recovery of *H*. *macrophthalmos* populations, further management actions such as temporary, spatial-temporal, or permanent closed seasons must be taken. In such a scenario, our SSPs for *H*. *macrophthalmos* would provide a powerful tool to detect the commercialization of illegal catches.

Groupers (subfamily Epinephelinae) and grunts (Haemulidae) represent two fish groups of high economic value and in high demand, which makes them major targets of substitution [38, 43, 46]. Herein we successfully developed SSP assays for one grouper (*Alphestes immaculatus*) and one grunt species (*Anisotremus interruptus*). Given that several species belonging to the subfamily Epinephelinae (e.g. *Alphestes* spp., *Epinephelus* spp., *Mycteroperca* spp.) are marketed simply as “*mero*”, our assay for the identification of *A. immaculatus* represents an important tool that can be used to identify processed forms of “*mero*” samples. In Peru, *A. interruptus* is referred to as “*burrito*” or “*chita dorada*” [41, 69] and more recently the common name “*berrugata*” has been attributed to this species in commercial venues from Piura region [44] (authors’ personal observation), which is also the market name of the Pacific tripletail *Lobotes pacifica* [41, 69]. The in-vitro validation results, which included fresh and cooked *A. interruptus* samples (electrophoresis gels A to E in Fig 12, Table 4) resulted in 100% accuracy and high specificity, without cross-reactions with the DNA from related and non-related species including *L. pacifica*. Thus, the *A. interruptus* identification assay presented herein will facilitate the rapid authentication of commercial samples labeled as “*berrugata*”.

### Conclusions, recommendations, and future perspectives

The seafood industry is growing globally and rapidly, increasing the need for effective fishery management measures and strong regulatory systems supported by modern and reliable molecular identification tools. We addressed the scarcity of species-specific identification assays for Peruvian seafood species by designing and validating novel SSPs targeting 10 important marine species from the Eastern Pacific. The robustness, versatility, and specificity of our novel SSP assays were demonstrated by the rapid and accurate identification of target species in marine eDNA and fresh and processed commercial samples. Our results included publically available primer sequences and complete validated molecular protocols that are ready to be used by regulatory or law enforcement bodies, research institutes, universities, and aquaculture corporations, providing support to seafood certification programs, future implementation of regulatory policies and traceability protocols, marine research, and aquaculture production.

We also presented the first species-specific eDNA assay developed in Peru for a marine species, which is a significant contribution to further population assessments of the highly economically important Peruvian scallop. Our eDNA assay was able to detect the presence of *A. purpuratus* in all sampling stations where this species occurs. Environmental DNA technique is an emergent technology used to estimate biomass or species abundance of several organisms, enhancing traditional methods employed during biomass estimates. Further efforts should be made to develop more eDNA assays for species detection and abundance estimation of priority Peruvian marine resources including protected, endangered, overexploited, highly commercial, and invasive species. In that sense, our novel SSPs targeting fish species can be of utility in the evaluation of further eDNA assays.

There is an urgent need to address the different issues caused by low taxonomic resolution during landings, mislabeling, illegal fishing, weak regulations, lack of traceability systems, and ineffective management of some Peruvian fishery resources. Initial efforts should be directed to standardizing the market names by the creation of an official list of unique commercial names, which was already recommended years ago [38, 43, 46] but not yet implemented. Also, further actions including population studies and the implementation of fishery regulations (e.g. permanent or reproductive closed seasons) must be taken to ensure the recovery of overexploited resources like *H. macrophthalmos*. Finally, the development of further molecular assays for seafood authentication and eDNA surveillance programs will significantly contribute to the efforts to mitigate overexploitation, illegal fishing, and mislabeling, leading to better management of our marine resources.

## Acknowledgments

We want to thank José Carranza (Nagoya University), Julissa Sánchez-Velásquez (University of Melbourne), and Diego Torres (National University of Trujillo) for their help during seafood sample collection. We also thank Percy Montero (Instituto del Mar del Perú) for his support during eDNA sampling in Tumbes region. We are also greatly in debt to anonymous fishermen, commercial divers, and fishmongers who sold or donated fish and shellfish samples toward this study.

## Supporting information

**S1 Table.** Fish and shellfish species used for non-cross species validation of the species-specific primers. GenBank and BOLD accessions are written between parenthesis and brackets respectively. Sampling venues correspond to FLS: fish landing site, MK: market, SMK: supermarket, and WFM: wholesale fish market

**S2 Table.** Primers used for barcoding identification of voucher specimens obtained in this study

**S1 Fig**. In-silico analysis of the binding sites of primer set ARGOF/ARPU129R and in *A. purpuratus* sequences (PDF)

**S2 Fig.** Direct qPCR and endpoint PCR assays for the detection of commercial samples of *Argopecten purpuratus*

**S3 Fig**. Multiplex PCR for the detection of cooked seafood samples: A) *Paralabrax callaensis* and *P. humeralis*, and B) *Hemilutjanus macrophthalmos* and *Schedophilus haedrichi*

**S4 Fig** In-silico analysis against non-target species of the species-specific primers for A: *Etropus ectenes*, B: *Hemilutjanus macrophthalmos*, C: *Schedophilus haedrichi*, D: *Alphestes immaculatus*, and E: *Anisotremus interruptus*

**S1 File**. Primer binding region of the SSP set for *Argopecten purpuratus* in all available DNA sequences of the target species in FASTA format (FASTA)

**S2 File**. Primer binding region of the SSP set for *Argopecten purpuratus* in 20 scallop species in FASTA format (FASTA)

**S3 File.** Primer binding region of the SSP set for *Doryteuthis gahi* in all available DNA sequences of the target species in FASTA format (FASTA)

**S4 File**. Primer binding region of the SSP set for *Doryteuthis gahi* in all available DNA sequences of non-target species *Dosidicus gigas* in FASTA format (FASTA)

**S5 File**. Primer binding region of the SSP set for *Paralichthys adspersus* in all available DNA sequences of the target species in FASTA format (FASTA)

S6 File. Primer binding region of the SSP set for *Paralichthys adspersus* in 10 congeneric species in FASTA format (FASTA)

**S7 File**. Primer binding region of the SSP set for *Etropus ectenes* in all available DNA sequences of the target species in FASTA format (FASTA)

**S8 File**. Primer binding region of the SSP set for *Paralabrax callaensis* in all available DNA sequences of the target species in FASTA format (FASTA)

**S9 File**. Primer binding region of the SSP set for *Paralabrax humeralis* in all available DNA sequences of the target species in FASTA format (FASTA)

**S10 File**. Primer binding region of the SSP set for *Hemilutjanus macrophthalmos* in all available DNA sequences of the target species in FASTA format (FASTA)

**S11 File**. Primer binding region of the SSP set for *Schedophilus haedrichi* in all available DNA sequences of the target species in FASTA format (FASTA)

**S12 File**. Primer binding region of the SSP set for *Alphestes immaculatus* in all available DNA sequences of the target species in FASTA format (FASTA)

**S13 File**. Primer binding region of the SSP set for *Anisotremus interruptus* in all available DNA sequences of the target species in FASTA format (FASTA)

